# Molecular formula discovery via bottom-up MS/MS interrogation

**DOI:** 10.1101/2022.08.03.502704

**Authors:** Shipei Xing, Sam Shen, Banghua Xu, Tao Huan

## Abstract

A substantial fraction of metabolic features remains undetermined in mass spectrometry (MS)-based metabolomics. Here we present bottom-up tandem MS (MS/MS) interrogation to illuminate the unidentified features via accurate molecular formula annotation. Our approach prioritizes MS/MS-explainable formula candidates, implements machine-learned ranking, and offers false discovery rate estimation. Compared to the existing MS1-initiated formula annotation, our approach shrinks the formula candidate space by 42.8% on average. The superior annotation accuracy of our bottom-up interrogation was demonstrated on reference MS/MS libraries and real metabolomics datasets. Applied on 155,321 annotated recurrent unidentified spectra (ARUS), our approach confidently annotated >5,000 novel molecular formulae unarchived in chemical databases. Beyond the level of individual metabolic features, we combined bottom-up MS/MS interrogation with global peak annotation. This approach reveals peak interrelationships, allowing the systematic annotation of 37 fatty acid amide molecules in human fecal data, among other applications. All bioinformatics pipelines are available in a standalone software, BUDDY (https://github.com/HuanLab/BUDDY/).

## Introduction

Tandem mass spectrometry (MS/MS) reveals the structural information of chemicals by measuring the mass-to-charge ratios (*m/z*) of ions fragmented from a selected precursor ion. MS/MS is essential in MS-based untargeted metabolomics for metabolite identification. A routine process is to match experimental MS/MS against reference MS/MS spectra^1–4^. However, given the great sensitivity of MS instruments, a considerable amount of uncharacterized chemical signatures (‘dark matter’) remain in untargeted metabolomics^5^. These unidentified features might present unique bioactivities and play critical roles in understanding biological mechanisms. Unfortunately, these features do not have available reference spectra in spectral databases or even have not been reported in literature (‘unknown unknowns’^6^). As such, unknown annotation has become a challenging yet pivotal research topic in metabolomics, exposomics, and others^1, 7–14^

Molecular formula determination is a great starting point to illuminate unrecognized metabolites, as unambiguous molecular formula annotation can dramatically reduce the number of potential chemical candidates. A common practice for molecular formula annotation relies on mass searching against metabolome databases (HMDB^15^, KEGG^16^, ChEBI^17^, etc.) or even larger chemical databases (PubChem^18^, ChemSpider^19^, etc.). However, this approach is significantly affected by the mass accuracy, limiting the searching scope within existing databases and thus hindering the discovery of novel formulae. Moreover, it does not fully utilize available information such as MS1 isotope pattern and MS/MS. SIRIUS first overcame the above issues by producing all mathematically possible formulae^20^, decomposing isotope patterns^21^, and integrating fragmentation trees^22^ into MS/MS analysis. SIRIUS works in a top-down manner in which formula candidates are first generated using MS1 information, and MS/MS explanation comes last.

Bottom-up processing represents specifying individual base components separately and assembling them into a top-level system. Bottom-up approaches have been widely adopted in neighboring fields such as shotgun sequencing^23, 24^ and shotgun proteomics^25, 26^, where DNA strands and intact proteins are sheared or digested into shorter sequences for more detailed analyses. Here we propose bottom-up MS/MS interrogation to enable accurate molecular formula determination with significance estimation^10, 27^. While a couple of bioinformatics tools have integrated MS/MS analysis into their candidate ranking^7, 9, 22, 28^ (formula or structure candidates), we first prioritize the significance of MS/MS in candidate generation. The ‘bottom-up’ methodological design dramatically shrinks the candidate searching space and allows for the broad discovery of unreported biochemically feasible formulae. Implementing machine-learned ranking (MLR) and false discovery rate (FDR) estimation enhances the annotation accuracy and enables significance control, respectively. Furthermore, we explored beyond the annotation of individual metabolic features up to the molecular network level—we present experiment-specific global peak annotation, which reserves reasonable individual peak annotations while revealing peak-peak relationships on an experimental basis.

## Results

BUDDY was created as a standalone platform with an intuitive graphical user interface (**Extended Data Fig. 1**), capitalizing on bottom-up MS/MS interrogation and experiment-specific global peak annotation for untargeted metabolomics. Conceptually, bottom-up MS/MS interrogation enables highly accurate molecular formula determination with significance estimation. On the other hand, experiment-specific global peak annotation aims to construct valid biotic or abiotic metabolic feature connections while refining individual feature annotations. A YouTube video is provided to enable a quick start guide (https://www.youtube.com/watch?v=Ne_Y0vZ0WKI).

### Bottom-up MS/MS interrogation

**Fig. 1** illustrates the schematic workflows of top-down and bottom-up approaches for molecular formula annotation. The top-down approach (e.g., SIRIUS^7^) generates the entire potential candidate space using MS1 information, assesses the MS/MS interpretability for each formula candidate (e.g., fragmentation tree^22^), and ranks them by heuristically combining MS1 score and MS/MS score. In comparison, our bottom-up approach has three distinct features, including (1) generation of an MS/MS-explainable candidate space, (2) machine learning-assisted candidate ranking, and (3) FDR estimation. Bottom-up MS/MS interrogation starts with decomposing the query MS/MS into multiple fragment-neutral loss pairs. The masses of each pair are searched against the well-curated formula database archiving >3.5 million unique molecular formulae from 26 chemical databases (**Methods**, **Supplementary Fig. 1**) to produce possible subformula candidates. Notably, the database search here does not limit the generation of molecular formulae from extending beyond the current chemical space but actually allows us to prioritize discovering unreported (bio)chemically feasible molecular formulae. During the search, both even-electron and odd-electron (radical) ion species are considered equally through searching against either the hydrogen-adjusted formula database (e.g., C_6_H_7_) or the original neutral molecular formula database (e.g., C_6_H_6_). This approach enables a broader recognition of radical ions, especially in collision-induced dissociation-based MS/MS spectra^29^ (**Methods**). Next, the subformula candidates for each fragment-neutral loss pair are stitched together to add a unique dimension to the formula candidate space. Candidate spaces of all dimensions are merged, dereplicated, and then subjected to SENIOR rules^30^ to filter out chemically implausible formulae (e.g., CH_5_). The above processes create a pooled MS/MS- explainable candidate space in which any candidate can explain at least one peak in the query MS/MS. MS1 isotope pattern matching is conducted optionally to generate MS1 isotope similarities. The following machine-learned ranking (MLR) assesses both intrinsic properties of candidate formulae and their performance in MS/MS interpretation, and an MLR prediction score is assigned to each candidate. MLR scores are then converted into estimated posterior probabilities using Platt calibration^31^, and FDR estimates are calculated accordingly (**Methods**).

**Fig. 1.**
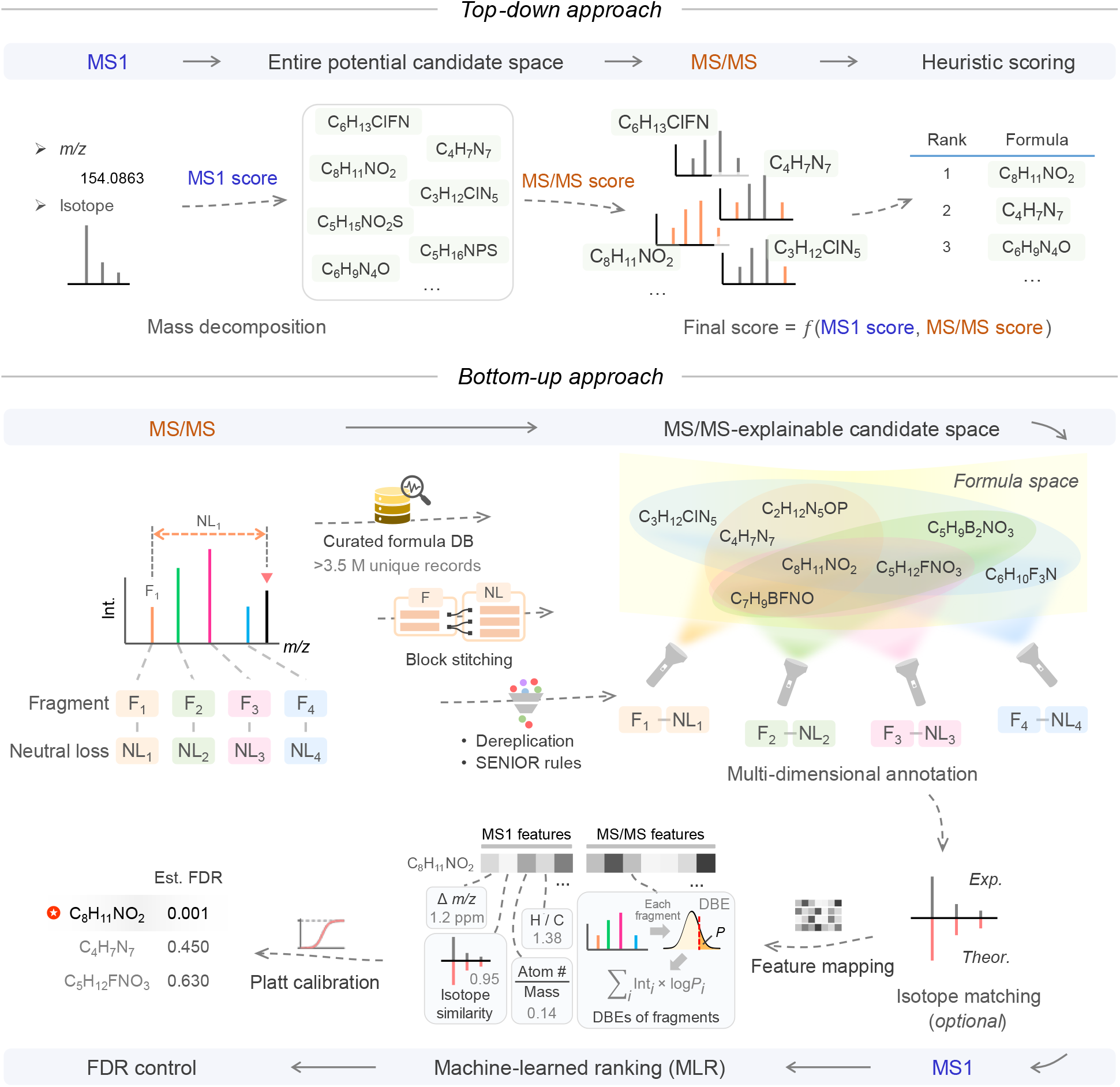
Methodological comparison between top-down and bottom-up approaches for molecular formula annotation. The top-down approach generates candidate formulae from MS1 information, followed by MS/MS explanation and candidate ranking. Heuristic scoring is applied to combine MS1 and MS/MS scores to calculate the final score. Bottom-up molecular formula determination prioritizes candidate formulae that can explain MS/MS in a chemically feasible manner. Multi-dimensional annotation drastically narrows down the candidate searching scope. Machine-learned ranking (MLR) is implemented for automated and accurate candidate ranking. False discovery rate (FDR) is controlled via Platt calibration.

### Candidate space shrinkage

The first key feature of the bottom-up approach lies in its capability to drastically shrink the candidate space for lower computational cost and higher annotation accuracy. To demonstrate this, we performed a repository-scale candidate space comparison between the top-down and bottom-up approaches using the curated NIST20 library (21,976 unique chemicals, **Methods**). For each query MS/MS, we generated both its entire potential candidate space and its MS/MS-explainable candidate space via mathematical mass decomposition^20^ and bottom-up MS/MS interrogation, respectively.

The results show that a narrower searching space can be obtained in 87.8% of total queries by prioritizing MS/MS-explainable candidates (**Fig. 2a**). Compared to the top-down approach, the bottom-up approach shrinks the candidate space by 4.2% to >99.9%, 42.8% on average (median of 40.8%). To further explore the candidate space shrinkage in the domain of precursor mass, we plotted the candidate counts of all queries in both spaces as in **Fig. 2b**. Overall, both candidate spaces expand as precursor mass increases, but the MS/MS-explainable candidate space grows at a drastically lower rate. As estimated by LOWESS, the MS/MS-explainable candidate space is 7.9-fold narrower than the entire space at *m/z* 400, 12.2-fold at *m/z* 600, and 20.7-fold at *m/z* 800. A more obvious size difference between the two spaces can be observed at higher *m/z.* Even for *m/z* <200, a statistical significance was found between the space sizes (*P* < 2.2×10^-16^).

**Fig. 2.**
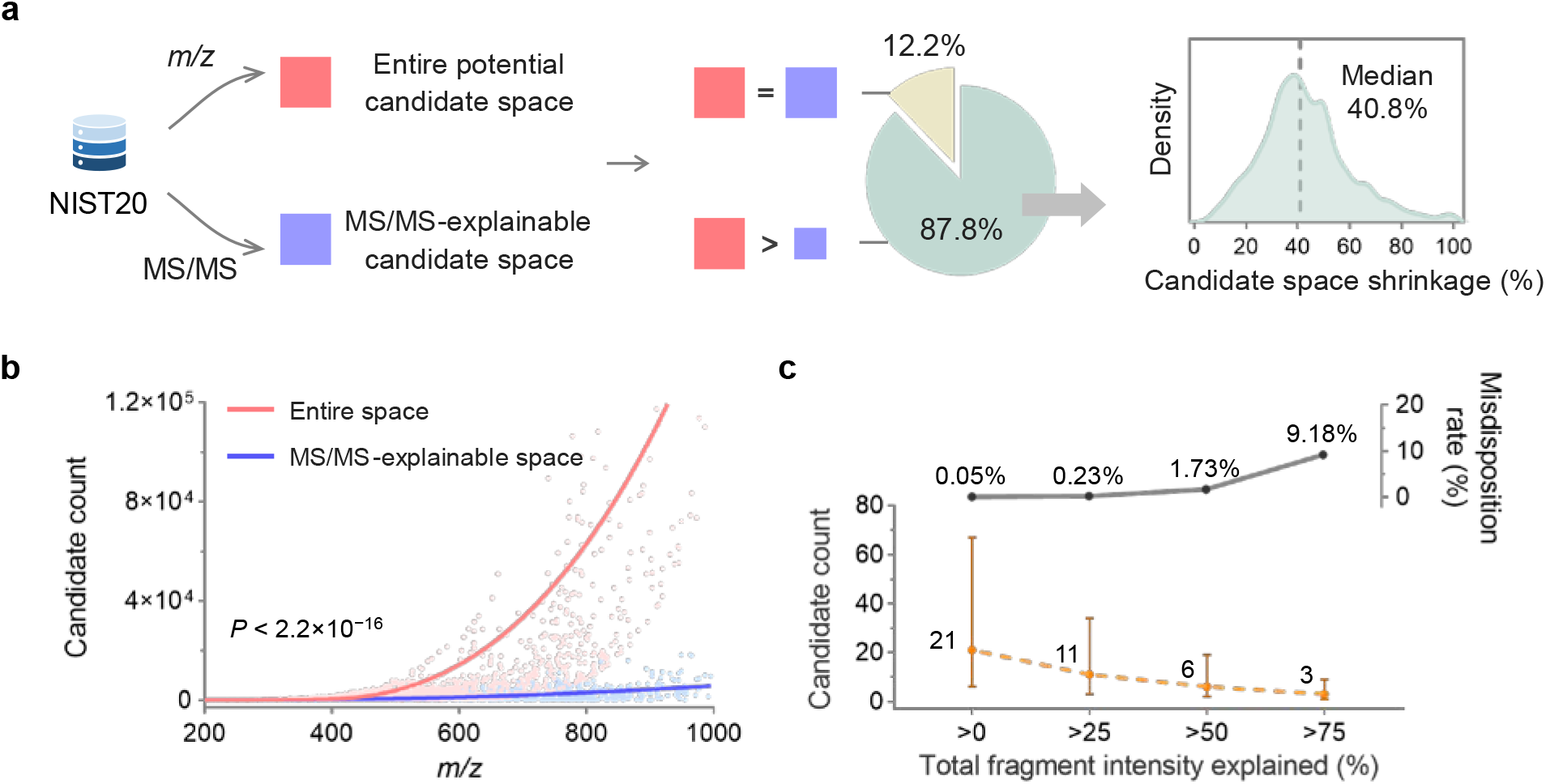
Bottom-up approach prioritizes MS/MS-explainable candidates. **a**, The MS/MS-explainable candidate space is narrower than the entire potential candidate space in 87.8% of the queries. **b**, The bottom-up approach provides a dramatically narrower candidate space, especially for large *m/z* values. Paired Mann-Whitney *U* test, two-sided, *P* < 2.2×10^-16^. **c**, A higher threshold of total explained fragment intensity further shrinks the candidate space (medians with interquartile ranges shown). The chance of disposing of correct answers (i.e., misdisposition rate) also increases with a higher cutoff but remains <2% when the cutoff of total explained fragment intensity is 50%.

Additionally, the MS/MS-explainable candidate space can be shrunk further by applying cutoffs on MS/MS explanation qualities, such as the total fragment intensity explained (normalized, in percentage). However, the process of candidate space shrinkage is always accompanied by the risk of mistakenly disposing of correct answers outside the candidate space, which we term misdisposition. This means that there are extreme cases where the correct formula cannot explain a single fragment-neutral loss pair in its MS/MS. The chance of misdisposition remains as low as 0.05% (1 out of 2000 annotations). With a higher threshold of the total fragment intensity explained, the candidate count reduces further at the cost of a higher misdisposition rate (**Fig. 2c**). In our bottom-up approach, we do not include any additional MS/MS-related cutoffs for candidate removal as experimental MS/MS can be informative to varying extents or even be contaminated due to collision energy or precursor window width selections^32, 33^. Hence, we prioritize a more accurate candidate ranking process while reserving a relatively reasonable fraction of formula candidates.

### Accuracy of bottom-up molecular formula annotation

The method performance was evaluated first on four reference MS/MS spectral libraries: MassBank, GNPS, Fiehn HILIC, and Vaniya-Fiehn Natural Products Library (VF-NPL). Data curation ensured structure-disjoint evaluation in each reference library (**Methods**). In total, 24,216 MS/MS spectra representing 11,667 unique molecules were reserved. These molecules cover 19 chemical superclasses and 263 classes^34^ (**Fig. 3a**, **Supplementary Table 1**), embracing a diverse range of structures. For method comparison, we chose SIRIUS^21^, a benchmarking tool for molecular formula determination. The same parameter set was applied in both tools. The results are summarized in **Fig. 3b** (spectra from Orbitraps) and **Extended Data Fig. 2** (spectra from QTOFs). Overall, BUDDY outperforms SIRIUS in all tested MS/MS libraries, improving the top 1 accuracy by an average of 30.1% (**Supplementary Table 2**); the largest accuracy increase was found in VF-NPL (Orbitrap, positive mode spectra) at 68.7%. We further investigated BUDDY’s performance from two aspects, precursor mass and spectral quality. As shown in **Fig. 3c**, the annotation accuracy remains high (93.0%) for *m/z* <400 (67.4% of total queries) but drops as precursor mass increases (63.8% accuracy for *m/z* ≥400). From the perspective of spectral quality, we herein used spectral entropy^35^ to refer to the extent of MS/MS fragmentation (chaos). In brief, a more informative MS/MS with more diverse fragments generally receives a higher spectral entropy and is thus anticipated to have a higher chance of being correctly annotated. We inspected annotation accuracy as a function of spectral entropy for compounds of *m/z* 300 to 400 and 400 to 500 (**Fig. 3d**) and found that while an increasing trend is observed for *m/z* 400 to 500 *(r* = 1.0), the relationship is unclear for *m/z* 300 to 400 (*r* = 0.2).

**Fig. 3.**
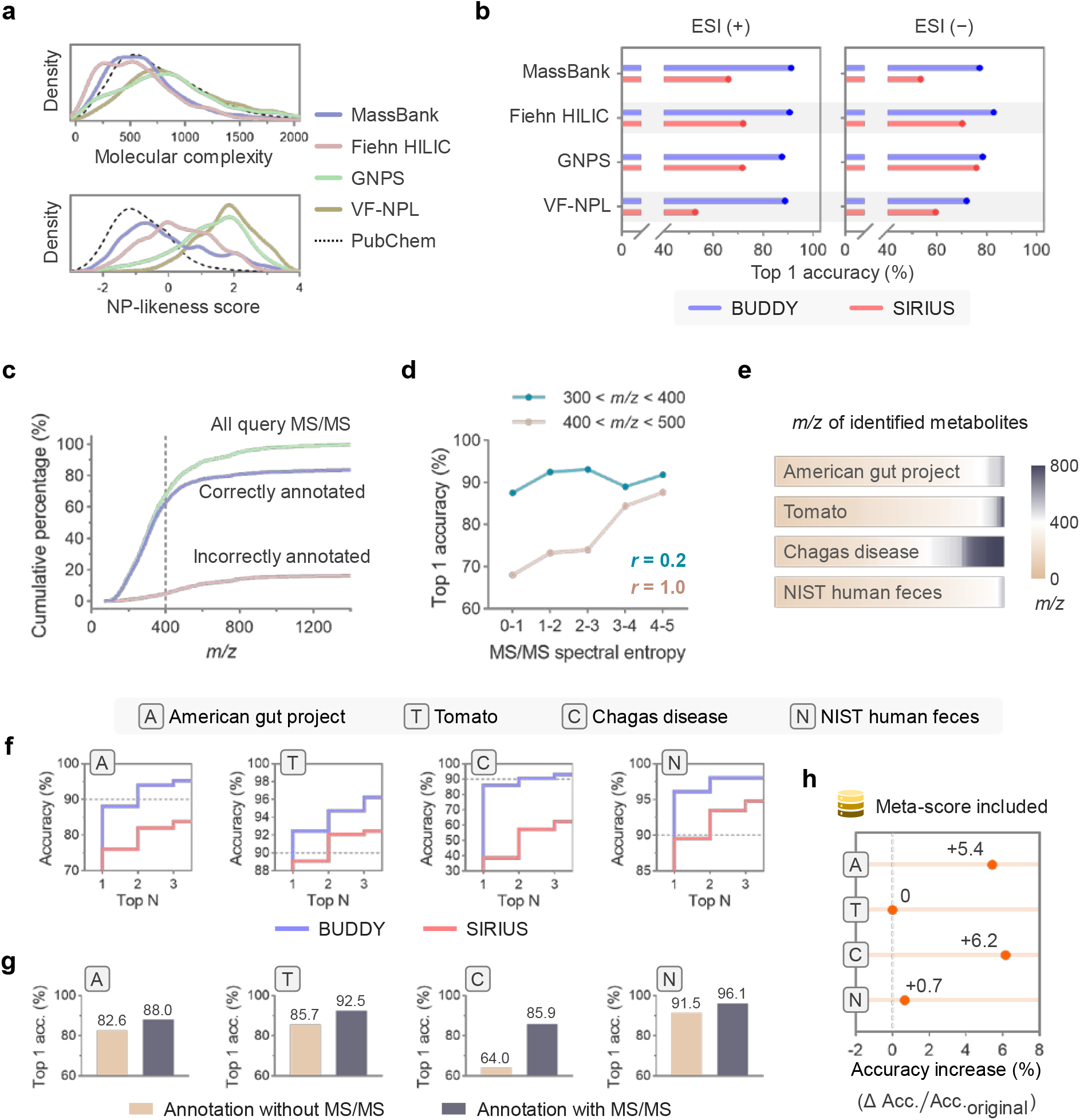
Method performance evaluation. **a**, Molecular complexity and natural product (NP)-likeness score of tested compounds in MS/MS spectral libraries and PubChem. Tested compounds cover a broad range of structural complexity. **b**, Method evaluation result on four reference MS/MS libraries (Orbitrap MS/MS spectra shown). ESI (+), electrospray positive ion mode; ESI (-), electrospray negative ion mode. **c**, Annotation performance on the *m/z* domain. **d**, Annotation accuracy of MS/MS with different spectral entropies (Spearman’s rank correlation coefficient shown). **e**, Mass distributions of identified metabolites in four public LC-MS/MS datasets. **f**, Annotation accuracy (top 1 to top 3) of BUDDY and SIRIUS on tested LC-MS/MS datasets. **g**, Annotation with MS/MS spectra provides higher accuracy in all tested datasets. **h**, Annotation accuracy increases (top 1) when meta-scores are included for formula determination.

Bottom-up molecular formula annotation was also evaluated on experimental data using four public LC-MS/MS-based metabolomics datasets collected from various sample matrices, MS instruments, and ionization polarities (**Supplementary Table 3**). The metabolites identified via spectral matching (level 2a^36^) were used as ground truths (**Fig. 3e**). Again, the bottom-up molecular formula annotation consistently outperforms the top-down approach in all datasets from top 1 to 3 accuracies (**Fig. 3f**, **Supplementary Tables 4-7**). The slightly worse performances in the American gut project (88.0%) and Chagas disease (85.9%) datasets could be respectively attributed to MS instrumental mass resolution (only the American gut project dataset was collected on QTOF and not Orbitrap) and precursor mass distribution (**Fig. 3e**). Additionally, we tested how MS/MS spectra aid in formula annotation and generated annotation results without using MS/MS (**Fig. 3g**). In all cases, the employment of MS/MS was positive, and the most significant improvement occurred in the Chagas disease dataset (34.2%). Moreover, BUDDY offers an option to use meta-scores^37^ (e.g., appearance in chemical databases) for the bottom-up annotation. In reanalyzing the above metabolomics datasets with meta-scores, an increased annotation accuracy (up to 6.2%) was observed in three of the datasets (**Fig. 3h**).

### Platt calibration and FDR estimation

MLR prediction scores are transformed into calibrated probabilities, on top of which FDR estimation can be achieved (**Methods**). Using MassBank^4^ as an initial validation, we explored MLR score distributions of both correct and incorrect annotations (**Fig. 4a**). Corresponding curves for calibrated probabilities show that 88.0% of correct annotations receive calibrated probabilities of >0.5, and 97.7% of incorrect annotations have calibrated probabilities of <0.5. The receiver operating characteristics (ROC) curve for differentiating correct and incorrect annotations (**Fig. 4b**) yields an area under the curve (AUC) of 0.985. Next, the performance of FDR estimation was assessed in all of the above-mentioned reference MS/MS libraries through Q-Q plots (**Fig. 4c-d**, **Extended Data Fig. 3**); the estimated FDR consistently shows correlations of >0.98 with exact FDR. However, note that FDRs are inclined to be overestimated to varying extents in real metabolomic datasets (**Supplementary Fig. 1**) due to MS/MS contamination.

**Fig. 4.**
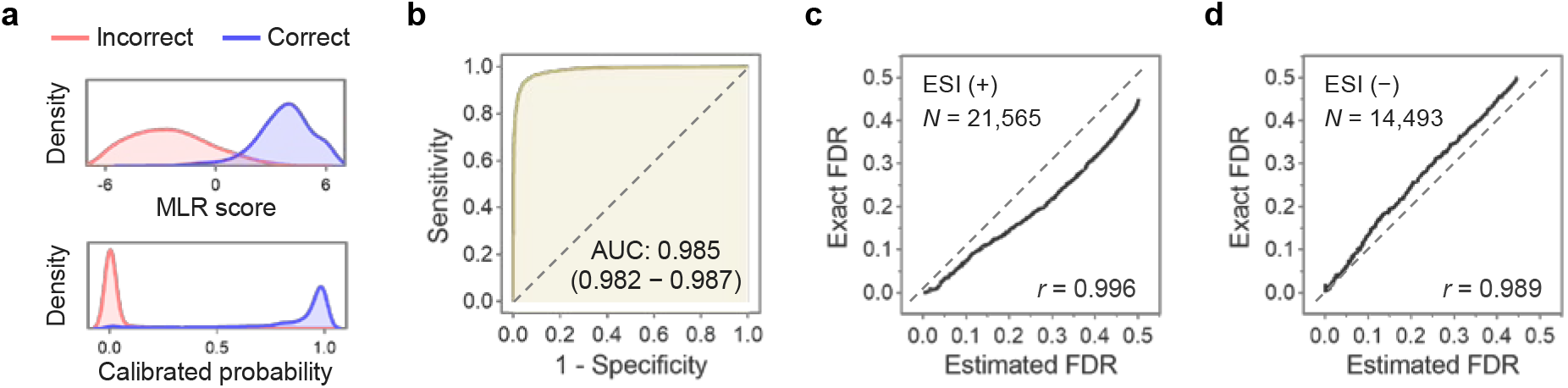
Platt calibration and FDR estimation. **a**, Density plots of MLR scores and calibrated probabilities for correct and incorrect annotations using MassBank (Orbitrap, positively ionized MS/MS). **b**, ROC plot classifying correct and incorrect annotations using calibrated probabilities. **c & d**, Evaluation of FDR estimation using Q-Q plots (Pearson’s correlation coefficients shown) in electrospray positively and negatively ionized spectra (MassBank).

### Bottom-up MS/MS interrogation for unreported formula annotation

As an initial test of bottom-up MS/MS interrogation to unravel unrecognized metabolites, we collected MS/MS spectra of five novel compounds discovered in references^10, 11^, three of which were confirmed by nuclear magnetic resonance (NMR) experiments. In all cases, BUDDY ranked the correct molecular formulae as top 1 (**Extended Data Fig. 4**), with the estimated FDR ranging from 0.6% (C_33_H_49_NO_5_, *m/z* 540.3690) to 16.3% (C_14_H_20_N_4_O_2_S, *m/z* 309.1383).

As a repository-scale application, we retrieved MS/MS libraries of annotated recurrent unidentified spectra (ARUS)^38^ collected from human plasma and urine samples. Data preprocessing led to 155,321 unidentified spectra (**Methods**). Overall, bottom-up MS/MS interrogation resolved 153,079 spectra (98.6%) and determined 5,191 formulae unarchived in HMDB or KEGG at <5% of the estimated FDR (**Fig. 5a**, **Supplementary Tables 8-11**). In particular, we discovered 173 and 53 completely novel molecular formulae absent from PubChem in plasma and urine, respectively. We manually inspected the MS/MS of a novel formula *(m/z* 674.4387, C_35_H_64_NO_9_P, **Fig. 5b**) and annotated its structure as PC(18:2/9:0(CHO)) using characteristic fragmentation patterns (**Supplementary Note 1**). Moreover, to demonstrate the biochemical feasibility of newly discovered formulae, we tried to connect them with existing formulae via common biochemical transformations (**Fig. 5c**, **Supplementary Table 12**). Transformations with up to two steps can link 95.4% (plasma) and 100% (urine) of novel formulae to known formulae (**Supplementary Table 13**). The remaining unlinked formulae are large-mass formulae with >60 carbons and could all be represented as homologues of known lipids with different acyl side chains.

**Fig. 5.**
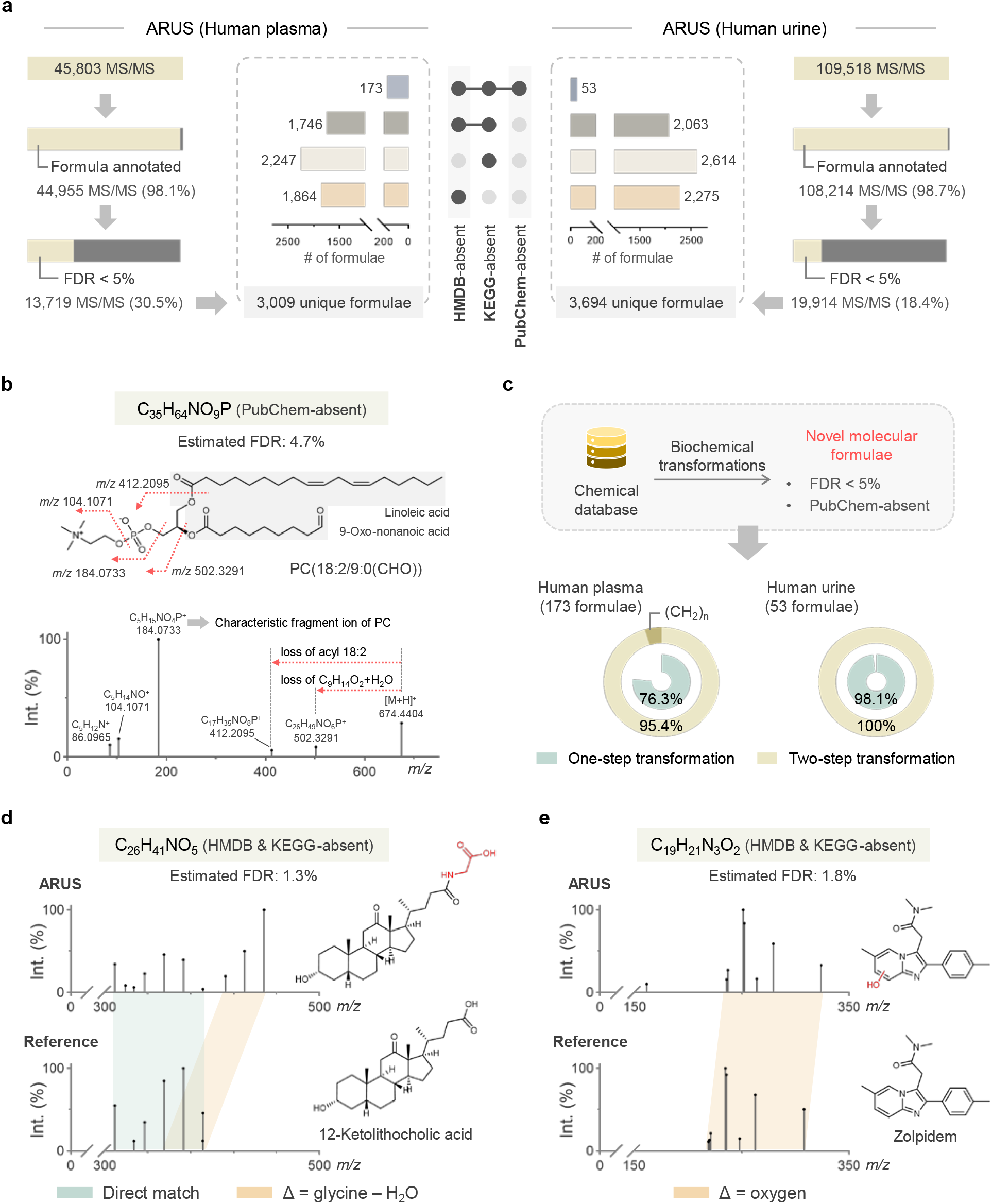
BUDDY discovers novel formulae absent from chemical databases. **a**, Formula annotation result summary of ARUS MS/MS libraries (human plasma and urine). **b**, Manual inspection of a novel formula absent from PubChem. **c**, Novel molecular formulae can be linked to existing molecular formulae via common biochemical transformations. **d** & **e**, Illustrations of orthogonal evaluation.

We additionally performed an orthogonal validation against MS/MS matching-based structural analog search (**Supplementary Note 2**), and consistent formula annotation results were obtained on 97.9% of tested spectra (**Supplementary Table 14**). We showcased an HMDB-absent annotation of glycine-conjugated 12-ketolithocholic acid (**Fig. 5d**). Searching this MS/MS against the entire public data repository using MASST^39^, this bile acid derivative was detected broadly in 28 metabolomic datasets collected from humans, mice, and bacteria. Another example of xenobiotic compounds detected in humans is depicted in **Fig. 5e**, where zolpidem is known as a hypnotic drug for patients with insomnia.

### Experiment-specific global peak annotation

Inspired by references^11, 13^, we extended the bottom-up MS/MS interrogation method beyond individual feature annotations up to the molecular network level to reveal peak-peak relationships on an experimental basis (**Methods**). In brief, the goal of global peak annotation is to select the optimal molecular network while considering both candidate posterior probabilities of individual features (node) and valid feature interrelationships (edge). This process updates the low-confidence peak annotations with the aid of MS/MS similarity-based peak mutual connections. Notably, BUDDY incorporates an additional step of experiment-specific mass deviation estimation to boost its annotation performance (**Supplementary Note 3**). We reanalyzed the tomato dataset (**Supplementary Fig. 2**) using three mass tolerances (5, 10, and 15 ppm). Even when the mass tolerance tripled to 15 ppm, the top 1 annotation accuracy only dropped by 0.4% to 93.2%. This distinct feature allows BUDDY to capture the actual mass deviations better and, on the other hand, provide more accurate FDR estimates.

### Application of BUDDY in untargeted metabolomics

As a complete application of BUDDY in untargeted metabolomics, we analyzed the NIST human fecal material standards dataset (MSV000086989), as shown in **Fig. 6a**. In total, out of 6,215 extracted metabolic features, 213 features (3.4%) were identified (level 2a) via an MS/MS library search against NIST20. BUDDY further annotated 5,733 unidentified features (92.2%); global peak annotation linked 5,092 annotated features (81.9%) to identified metabolites directly or indirectly, paving the way for downstream annotation propagation. At <5% estimated FDR, we discovered 134 formulae absent from HMDB and 215 formulae absent from KEGG (**Extended Data Fig. 5**).

**Fig. 6.**
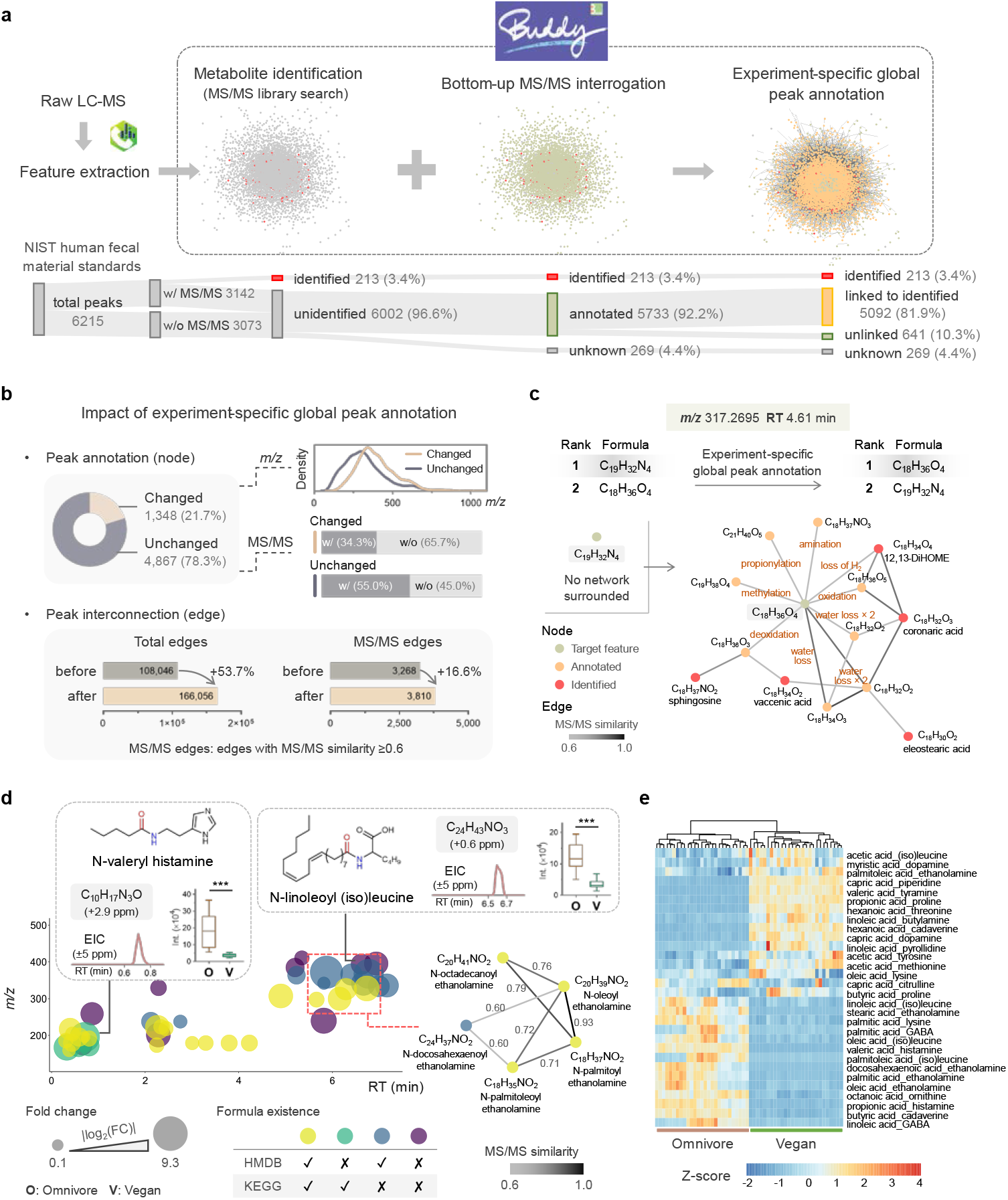
Experiment-specific global peak annotation and its application in untargeted metabolomics. **a**, Application of BUDDY on NIST human fecal material standards. In total, 81.9% of extracted metabolic features can be linked to identified metabolites directly or indirectly. **b**, Impact of experiment-specific global peak annotation in changing individual peak annotations and constructing metabolic feature interconnections. **c**, An illustrative example of experiment-specific global peak annotation. The biochemical transformations surrounding the target metabolic feature are labeled. Only MS/MS edges are shown. **d**, The bubble plot of FAAs annotated in human fecal material standards. ***: *P* < 1×10^-9^ (paired Mann-Whitney *U* test, two-sided, corrected by the Benjamini-Hochberg method). **e**, A heatmap showing FAA level differences between omnivores and vegans. FAA names are displayed in “fatty acid_amine” format. FAAs with fold changes >1.2 and adjusted *P* values <1 × 10^-3^ are included.

Next, we investigated the impact of global peak annotation from two aspects, individual peak annotations (node) and peak interconnections (edge). The global peak annotation process updated individual peak annotations for 1,348 (21.7%) features (**Fig. 6b**). Further analyses indicate that the annotation-changed peaks have comparably larger *m/z* values than unchanged peaks. Also, there were more MS/MS-unassigned peaks (65.7%) in annotation-changed peaks than in unchanged peaks (45.0%). This agrees with our intuition that higher *m/z* features without MS/MS spectra tend to be annotated with low confidence and are thus prone to be corrected in global peak annotation. On the other hand, global peak annotation increased the total count of peak interconnections by 53.7%. Focusing on the edges with MS/MS similarity ≥0.6 (MS/MS edges), 16.6% more MS/MS edges were constructed with updated formula annotations. We specifically showcased one annotation-changed peak in **Fig. 6c**. After global peak annotation, this peak was successfully connected to five identified metabolites and eight formula-annotated peaks via MS/MS edges. Corresponding biochemical transformations further reveal the structural insight, a dihydroxylated long-chain fatty acid, supported by subformula annotations of MS/MS fragments (**Supplementary Fig. 3**).

Finally, we systematically explored novel fatty acid amide (FAA) molecules in human feces. FAAs are essential signaling compounds that can be synthesized both endogenously and by the gut microbiota. Bacteria in the gut microbiota produce structural mimics of endogenous FAAs to interact with human cellular targets and thus modulate their hosts^40^. We generated 1,325 plausible FAAs using the fatty acids and amines present in the human gut^41^ and searched their molecular formulae against BUDDY annotations. Putative FAAs were first refined via characteristic fragmentation patterns. Further confirmation was performed by matching MS/MS against reference MS/MS of the corresponding fatty acids or amines using the reverse dot product (**Supplementary Table 15**). This exploration resulted in 37 annotated FAA molecules (**Fig. 6d**), and nine were molecular formulae absent from HMDB and KEGG. We manually investigated the feature annotated as N-valeryl histamine, which has an MS/MS spectrum that shows a reverse dot product of >0.99 with histamine (**Extended Data Fig. 5**). The local molecular network surrounding N-oleoyl ethanolamine (**Fig. 6d**) helps further unravel other N-acyl ethanolamine analogs through MS/MS similarities and biochemical transformations. Using a fold change cutoff (>1.2) and statistical analysis (adjusted *P* < 1×10^-3^), thirty FAA molecules were found to be significantly altered between omnivores and vegans (**Fig. 6e**), showing great potential for revealing the mechanisms of host-gut microbiota interactions and inter-individual differences in dietary response from the perspective of small molecules.

## Discussion

In bottom-up processing, individual base blocks are specified and assembled into a top-level system. Herein, we propose an MS/MS interpretation strategy in a bottom-up fashion, where each fragment ion and neutral loss pair offers a unique dimension to aid formula annotation. In the bottom-up approach, the MS/MS-explainable candidate space could be substantially narrower than the entire candidate space (e.g., ~20 folds at *m/z* 800). This design considerably reduces the computational cost for large-mass features from hours to seconds per query. More importantly, incorporating MLR and FDR estimation significantly improves the performance of bottom-up interrogation compared to the top-down approach (up to 68.7% accuracy increase) in uncovering both known and unknown molecular formulae while informing estimates of annotation confidence.

A more distinguished advantage of the bottom-up strategy is that it allows novel formulae exploration beyond the current chemical space while focusing on biochemically feasible ones. Although metabolome databases such as HMDB and KEGG often serve as reference repositories for biochemical studies, we confidently discovered >5,000 database-unarchived molecular formulae in human samples and implied their potential biological interests. These novel formulae could also complement molecular-formula-oriented data acquisition approaches, such as HERMES^42^. At the same time, it remains controversial whether to integrate meta-scores (e.g., citation frequency) in unraveling metabolic identities or not; meta-scores can help annotate known molecules^37^ at the cost of potentially discarding novel metabolites^10^. To this end, we offer an option of meta-score inclusion in BUDDY.

Our MLR module is dedicatedly designed to automatically integrate MS1 and MS/MS scores and rank formula candidates. One challenge is that the training data are insufficient to cover the entire range of MLR features. For example, the maximum double-bond equivalent (DBE) value within the training data is 40, and there could be unreported formulae with DBE >40. To allow the discovery of completely novel formulae, we mapped each MLR feature onto a distribution curve and used the two-sided *P* value for MLR training (**Methods**). Other careful considerations went into mimicking experimental conditions and improving the generalization of trained models, including precursor *m/z* shifting and MS1 isotope simulation (**Methods**). Last but not least, data augmentation employed in training MS/MS spectra further enhances the model robustness.

Certainly, MS/MS spectral quality is essential for annotation performance. Contamination fragments diminish the annotation confidence by lowering MS/MS explanation coverage or, even worse, generating incorrect candidate formulae that misguide subsequent interpretation. In this regard, annotation of unidentified features may benefit from targeted MS/MS profiling^42^ or more advanced data-driven spectral deconvolution approaches^43–45^. We also highly recommend that metabolomics practitioners consider mixed or ramped collision energies to collect information-rich experimental MS/MS in an untargeted manner.

Efforts have been made to control FDR on MS/MS spectral matching-based metabolite identification^27, 35, 46^, yet the FDR estimation for unknown annotation^10^ remains elusive. In our approach, Platt scaling monotonically maps MLR prediction scores onto a probability scale. However, note that accurate FDR estimates are expected only when experimental data quality is comparable to that of reference data. This means that both explicit MS1 isotope patterns and informative MS/MS spectra are responsible for reliable FDR estimates. We argue that accurate FDR control in untargeted metabolomics is still in its infancy, and both heuristic insights and statistical approaches should be involved.

Furthermore, we integrate bottom-up MS/MS interrogation with global optimization, aiming to synergistically resolve all the individual metabolic features on an experimental basis. For a more comprehensive and confident global annotation in untargeted metabolomics, combining orthogonal information, such as retention time^47–49^ and collision cross-section^50^, will be considered for future improvements. Meanwhile, molecular networking should embrace a variety of MS/MS similarity measures^8, 51–53^ for structural analog search to construct faithful connections among MS/MS spectra. Finally, in the pursuit of unambiguous structure annotation, the next generation of global networking may rely on a more convincing level of molecular substructures^54^ beyond molecular formula. With the advent of cheminformatics tools for in silico generation of hypothetical metabolites^55, 56^, we anticipate more structural insights into the undiscovered biosphere in recognition of a broad scope of unreported small molecules and their biological functions.

## Methods

### BUDDY software

BUDDY can perform three major tasks: (1) MS/MS library search, (2) bottom-up MS/MS interrogation, and (3) experiment-specific global peak annotation. For MS/MS library search, BUDDY offers three MS/MS matching algorithms: dot product, reverse dot product, and spectral entropy similarity. Users can upload and search against any MS/MS spectral library in MSP format. For bottom-up MS/MS interrogation, automated MS/MS preprocessing procedures are provided, including MS/MS noise peak elimination (**Supplementary Note 4**) and MS/MS deisotoping (**Supplementary Note 5**). Other settings regarding formula annotation include elemental restriction, chemical database restriction, elemental ratio restriction^57^, etc. Experiment-specific global peak annotation is modeled as an integer linear programming (ILP) problem, and it is a parameter-free module. BUDDY supports the import of both single and batch queries (metabolic feature table, MGF, or mzML file).

On the technical aspect, BUDDY was written in C# on the Universal Windows Platform. To speed up computation, BUDDY automatically invokes multiple processing cores and performs parallel programming. More details can be found in the user manual (https://github.com/HuanLab/BUDDY).

### Formula database construction and curation

To construct the neutral formula database, a total of 26 chemical repositories were downloaded, combined, and curated. These chemical repositories cover small molecules of metabolites, lipids, natural products, xenobiotics, drugs, toxins, contaminants, and so on (**Supplementary Fig. 4, Supplementary Table 16**). Currently, the neutral formula database embedded in BUDDY has 3,514,066 unique valid molecular formula records in total, incorporating ANPDB (2,157), BLEXP (27,531), BMDB (3,577), ChEBI (18,044), COCONUT (64,404), DrugBank (8,508), DSSTOX (150,250), ECMDB (1,528), FooDB (8,310), HMDB (10,968), HSDB (3,425), KEGG (8,550), LMSD (7,397), MaConDa (254), MarkerDB (561), MCDB (688), NORMAN (40,322), NPASS (10,009), Plantcyc (1,685), PubChem (3,507,371), SMPDB (2,467), STOFF-IDENT (7,753), T3DB (827), TTD (2,303), UNPD (28,899), and YMDB (1,060); the unique molecular formula record count in each repository is shown in parentheses. All of the above-mentioned chemical repositories were downloaded after December 2020. For each repository, we retrieved the following chemical information if provided: chemical name, molecular formula, CAS number, PubChem CID, KEGG ID, HMDB ID, SMILES, InChI, and InChIKey. The PubChem Identifier Exchange Service (https://pubchem.ncbi.nlm.nih.gov/idexchange) was used for conversion among various chemical identifiers. Chemical elements were restricted to C, H, N, O, P, S, F, Cl, Br, I, Si, B, Se, Na, and K. Charged and radical formulae were neutralized by a proton or hydrogen adjustment. Formulae with double-bond equivalent (DBE) values ≥-5 and monoisotopic masses ≤1500 were reserved. All formula strings were normalized using the Hill system and dereplicated. The final neutral formula database serves as (1) the original formula searching database, (2) the candidate formula database when database restriction is applied, and (3) the formula database for distribution analyses assisting MLR feature generation (see below).

### Radical fragmention

An important feature of BUDDY lies in its automatic recognition of radical fragment ions in MS/MS spectra. BUDDY shows no bias in recognizing odd-electron (radical) fragment ions and even-electron fragment ions. Similar approaches are also applied to neutral loss searching. Details are discussed in **Supplementary Note 6**.

### Adduct form

BUDDY allows numerous adduct forms for both ion modes, including [M + H]^+^, [M + NH_4_]^+^, [M + H – H_2_O]^+^, [M + Na]^+^, [M + K]^+^, [M – H]^-^, [M + Cl]^-^, [M + Br]^-^, [M + HCOOH – H]^-^, [M + CH_3_COOH – H]^-^, etc. When importing MGF files or metabolic feature tables into BUDDY, predefined adduct forms in the imported files can be automatically loaded and synchronized for downstream analysis. Users can change the adduct form for each MS/MS spectrum in the BUDDY graphical user interface. Users are also allowed to customize new adduct forms. Specifically, for adducts containing sodium or potassium atoms (e.g., [M + Na]^+^, [M + K]^+^, [M + Na – 2H]^-^), BUDDY also considers sodiated and potassiated fragment ions in addition to (de)protonated fragment ions and radical fragment ions^58^.

### MS1 isotope similarity

To compare experimental MS1 isotopes and theoretical isotopes of formula candidates, the isotope similarity calculation is implemented in BUDDY. Here we adopted and modified the isotope similarity algorithm from ref. ^28^ (**Supplementary Fig. 5**) for its broad scalability and low computational cost. We tested the validity of the isotope similarity algorithm (**Supplementary Note 7**, **Supplementary Fig. 6**), and the results show that experimental isotopes have statistically significantly higher similarity scores with theoretical isotopes of ground-truth molecular formulae than other formulae generated within a 5-ppm mass tolerance.

### Machine-learned ranking (MLR)

In BUDDY, formula candidates are ranked automatically through MLR. Here we describe the MLR feature design, training data preparation, data augmentation, model training, and hyperparameter optimization.

#### MLRfeature design

Both MS 1 – and MS/MS-related features were incorporated into the MLR task. In brief, MS1-related MLR features reflect the degree of similarity between experimental MS1 data and theoretical values; the features include *m/z* deviation, MS1 isotope similarity, and the intrinsic properties of candidate formulae (e.g., DBE and hydrogen-to-carbon ratio). MS/MS-related features indicate the performance of candidate formulae in explaining MS/MS spectra; these features include total intensities of explained fragments, DBEs of explained fragments, etc. Currently, a total of 38 MLR features are included in BUDDY, 14 of which are MS1-related and 24 are MS/MS-related. A complete list of MLR features and their descriptions are shown in **Supplementary Table 17**. Noting that the training data are insufficient to cover the entire range of MLR features, we thus mapped each MLR feature onto a distribution curve and used the two-sided *P* value for MLR training (**Supplementary Note 8**). We systematically analyzed the large-scale formula (MS1 -related MLR features) and NIST20 (MS/MS-related MLR features) databases and fit all distribution curves individually with skew normal distribution. Relevant results are shown in **Supplementary Tables 18-19** and **Supplementary Figs. 7-8**.

#### Training data preparation

NIST20 was purchased from the National Institute of Standards and Technology (NIST) through Isomass Scientific Inc. The high-resolution NIST20 database contains 1, 026,717 MS/MS spectra for 27,613 unique compounds. For model training purposes, NIST20 was curated as follows. First, we removed MS/MS spectra with isotopic adducts (e.g., MS/MS spectra for M + 1 ions), multiply charged adducts, or adducts containing chemical elements beyond the aforementioned elemental list (e.g., [M + Li]^+^). MS/MS spectra without InChIKey information were also discarded. To ensure structure-disjoint training, we performed spectral merging next (see references^7, 9, 13^ for the importance of structure-disjoint training). For each unique compound in NIST20 (identified by InChIKey), we chose its most frequent adduct form and performed spectra merging for MS/MS collected under different collision energies. Both fragment *m/z* values and ion intensities were averaged. The above curation processes led to 25,456 positively ionized and 11, 583 negatively ionized unique chemicals in NIST20.

Another training data preparation step was precursor *m/z* shifting. All precursor *m/z* values provided in NIST20 are recalibrated theoretical values and cannot be directly used for model training; the resulting trained models would be highly dependent on the MLR feature ‘precursor *m/z* error’. An intuitive explanation is that the candidate formula with the smallest absolute precursor *m/z* error (an absolute *m/z* error of 0) will always be the correct answer. Hence, we shifted the theoretical precursor *m/z* values within a specific range for experimental simulation. As described in ref. ^21^, mass deviations usually conform to the Gaussian distribution with a standard deviation that is 1/3 of the relative mass tolerance. However, we found lower real-case mass deviations, where the standard deviations of Gaussian curves are closer to 1/5 of the common relative mass tolerance for high-resolution MS data. We thus randomly sampled mass deviations from Gaussian curves within the mass tolerance range and shifted precursor *m/z* values from their theoretical values by the preselected mass deviations. The following mass tolerances were used: Orbitrap, 5 ppm; QTOF, 10 ppm.

Similarly, MS1 isotope simulation was performed to mimic the experimental conditions. We designed an MS1 isotope simulation workflow and validated its credibility (**Supplementary Note 9**, **Supplementary Figs. 9-10**, **Supplementary Table 20**). These simulated isotopic patterns were later compared with theoretical isotopic patterns of candidate formulae to compute the isotope similarity scores used for model training.

#### Data augmentation

As MS/MS spectra from multiple collision energies in NIST20 were merged for unique chemical compounds before training, there can be many more fragments in the merged MS/MS than the experimental MS/MS. Training data were thus augmented to enhance the robustness and generalization of trained MLR models (**Supplementary Note 10**).

#### Model training and hyperparameter optimization

MLR models were trained using LightGBM in the ML.NET framework for its superior performance on the structured data. Both data subsampling and MLR feature sub-sampling were applied to avoid overfitting. Data sub-sampling fraction was set as 0.9 and performed every 5 iterations. MLR feature sub-sampling fraction was set as 0.8. We set the learning rate as 0.005, and normalized discounted cumulative gain (NDCG) was used as the evaluation metric (**Supplementary Note 11**). Hyperparameters were optimized on the validation dataset (20% of the total training data) via grid search. These hyperparameters include L2 regularization term weight, number of iterations, maximum bin count per feature, and maximum number of leaves in one tree (**Supplementary Table 21**).

### Platt calibration and false discovery rate (FDR) estimation

In classification problems, probability calibration helps scale the distorted class probability distributions predicted by various classification models so that uninterpretable model prediction scores can be converted into corresponding class probabilities. Similarly, our MLR task can be simplified as a binary classification problem to some extent by dividing all the candidate formulae into two groups, correct and incorrect answers. Thus, probability calibration can also be applied to an MLR score to estimate its posterior probability *P(correct\MLR score),* the probability that a candidate is correct given the MLR prediction score. Platt calibration^31^ (or Platt scaling) is a common approach for probability calibration that learns a logistic regression model which maps scores *X* ∈ ℝ onto a scale of *P* ∈ [0,1]. We completed Platt calibration using the Platt calibrator of ML.NET in C#.

Furthermore, inspired by references^10, 27^, we estimated the FDR using posterior probabilities obtained by Platt calibration. FDR is defined as

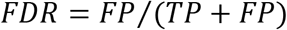

where TP and FP stand for true positives and false positives, respectively. Given a score threshold *x*_0_, FDR reflects the percentage of correct hits among all the hits receiving scores ≥ *x*_0_. In our case, considering the top *N* hits for a query metabolic feature that are associated with a certain score threshold, the total number of TP and FP is *N* (*TP* + *FP* = *N*). As the posterior probability can be interpreted as the expected value for a hit to be correct, we can estimate the number of TP within top *N* hits as the sum of their posterior probabilities 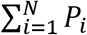. FDR can thus be estimated as

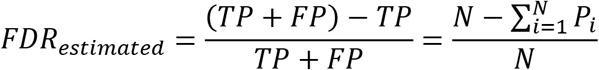

In practice, both calibrated Platt probability and estimated FDR are output by BUDDY for each hit so users can export batch results and easily apply FDR control to focus on metabolic features with high-confidence predictions. However, we must point out that FDRs can be overestimated due to various reasons, including but not limited to chimeric experimental MS/MS spectra, insufficient MS/MS fragments for annotation, and coeluting peaks that affect MS1 isotope information.

### Experiment-specific global peak annotation

BUDDY incorporates a global optimization approach for peak annotation corrections within the same LC-MS experiment, aiming to reveal peak-peak relationships while reserving reasonable individual peak annotations^11, 13^. Here a molecular network is constructed, where each peak is a node and every valid peak connection is an edge. Global peak annotation selects the optimal network considering both node scores and edge scores, and then some peak annotations generated by bottom-up MS/MS interrogation are updated. Notably, global peak annotation is completely free of meta-scores. Global optimization is achieved through integer linear programming via the OR-TOOLs package developed by Google. Linear constraints are set to guarantee a self-consistent network. More details are discussed in **Supplementary Note 12**. Tested on the NIST human feces dataset (6,215 peaks), global peak annotation took about 3 min on a personal computer (Intel i7-8700K CPU @ 3.70 GHz, Windows 10 64-bit operation system, 6 cores, 32 GB RAM).

### Candidate space shrinkage analysis

For candidate space shrinkage analysis, we used the curated positively ionized NIST20 library. MS/MS spectra with precursor masses larger than 1000 Da were removed, as the entire candidate space generation for large-mass chemicals can be extremely computationally expensive (up to hours or even days for a single query). The chemical element set was restricted to CHNOPSFClBrI. We retained MS/MS spectra of [M + H]^+^. This resulted in 21,976 unique chemicals for candidate space shrinkage analysis. In both candidate spaces, formula candidates within 5 ppm of the query precursor mass were retrieved to represent analysis results from high-resolution MS. Trendlines in **Fig. 2b** were generated using locally weighted scatterplot smoothing (LOWESS).

### Evaluation datasets

BUDDY was evaluated on diverse datasets, including publicly available reference MS/MS libraries, real metabolomics datasets, and ARUS MS/MS libraries.

#### Publicly available MS/MS libraries

Four public MS/MS libraries were downloaded in August 2021, including MassBank, GNPS, Fiehn HILIC, and Vaniya-Fiehn Natural Products Library (VF-NPL). From these libraries, we selected the reference MS/MS spectra with intact information for MS instrument type, InChIKey, molecular formula, and precursor adduct type. MS/MS spectra of adduct forms [M + H]^+^ or [M – H]^-^ were reserved. MS/MS spectra with precursor *m/z* deviations beyond the mass tolerance range were discarded. Furthermore, given the precursor mass (*m/z*)_pre_, we removed fragment ions with masses above (*m/z*)_pre_ – 0.5 in each MS/MS spectrum to ensure precursor ion exclusion. We categorized the reference spectra of each MS/MS library into batches according to their MS instrument type and ion mode. Within each batch, we kept unique chemical compounds (identified by InChIKey) for structure-disjoint evaluation. Well-curated MS/MS libraries were written into separate MGF files that can be directly imported into BUDDY or SIRIUS (settings in **Supplementary Note 13**). Evaluation results are summarized in **Supplementary Table 2**. For chemical structural diversity analysis, the molecular complexity^59^ and natural product (NP)-likeness score^60^ were computed in Python using RDKit, an open-source cheminformatics software. All parameters were set as default. We randomly selected one million molecules in PubChem for computation.

#### Public LC-MS/MS datasets

We downloaded four metabolomics datasets on MassIVE, covering diverse sample types collected from different MS instruments in positive or negative ion modes (**Supplementary Table 3**). Data preprocessing was conducted as follows. Raw LC-MS/MS data were converted into mzML files using MSConvert by ProteoWizard^61^, with all converted files in the centroided mode. MS-DIAL^44^ (version 4.70) was used to perform metabolic feature extraction and alignment, and aligned feature tables containing MS1 isotopes and MS/MS spectra were exported (see detailed parameters in **Supplementary Note 14**). To build up ground truths for formula determination, we conducted metabolite identification by searching experimental MS/MS against NIST20 using dot product^62^. We set the dot product score threshold as 0.7 and the minimum number of shared peaks (other than precursor ions) as 6 for high-confidence identification^27^. For repeatedly identified metabolites, we only reserved the feature with the highest dot product score (detailed settings are in **Supplementary Note 15**). This led to a total of 700 identified metabolites (level 2a^36^) in four datasets. Specifically, in the Chagas disease dataset (murine large intestine samples), more large-mass lipid molecules were identified *(m/z* >400 for 33.3% of identified compounds), which made it more challenging.

For the application of BUDDY, we used NIST human fecal material standards (MSV000086989). Peak extraction was completed in MS-DIAL. We then performed blank removal by discarding features with average intensities lower than twice that of the method blank. The remaining features were imported into BUDDY for metabolite identification, bottom-up MS/MS interrogation, and global peak annotation (settings in **Supplementary Note 16**). MS/MS spectra of putative fatty acid amides were compared against the reference MS/MS of both their corresponding fatty acids and amines using the reverse dot product, and we reserved the features with a similarity score >0.7 in either comparison.

#### ARUS MS/MS libraries

Annotated recurrent unidentified spectra (ARUS)^38^ libraries archive recurring MS/MS spectra of unknown identity in specific biological samples. Currently available ARUS libraries were collected from human plasma and urine using Orbitraps in both ionization polarities (https://chemdata.nist.gov/dokuwiki/doku.php?id=chemdata:arus). We carried out spectral clustering for redundancy removal. Similar MS/MS spectra were identified by precursor mass match (5 ppm) and dot product score (≥0.95). For MS/MS spectra of the same identity, we reserved the spectrum with the most fragments. This led to a total of 45,803 unidentified spectra from plasma and 109,518 spectra from urine. Detailed settings in BUDDY can be found in **Supplementary Note 17**.

### SIRIUS analysis

SIRIUS^7^ (version 4.9.2) was downloaded from https://bio.informatik.uni-jena.de/software/sirius. For molecular formula determination, we set the timeout for a single molecular formula (‘Tree timeout’) as 50 sec and the timeout for an MS/MS spectrum (‘Compound timeout’) as 200 sec. All SIRIUS analyses were conducted on an Intel i7-8700K CPU @ 3.70 GHz with 6 cores and 32 GB memory (Windows 10, 64-bit operating system).

## Data availability

The four reference MS/MS libraries used for evaluation can be freely downloaded at https://mona.fiehnlab.ucdavis.edu/downloads. ARUS MS/MS libraries can be downloaded at https://chemdata.nist.gov/dokuwiki/doku.php?id=chemdata:arus. All LC-MS/MS datasets are available from the MassIVE repository (American gut project, MSV000081981; tomato, MSV000081463; Chagas disease, MSV000086988; NIST human fecal material standards, MSV000086988 and MSV000086989). Evaluation results are provided in supplementary tables.

## Code availability

BUDDY is written in the C# language on the Universal Windows Platform (UWP). It currently works in the Windows OS (Windows 10 or higher). The standalone software can be freely downloaded from GitHub (https://github.com/HuanLab/BUDDY/releases/tag/v1.0). Source codes are also available on GitHub (https://github.com/HuanLab/BUDDY) under the MIT License.

## Acknowledgments

This study was funded by the University of British Columbia Start-up Grant (F18-03001), Canadian Foundation for Innovation (CFI 38159), and Natural Sciences and Engineering Research Council of Canada (NSERC) Discovery Grant (RGPIN-2020-04895). We thank Alisa Hui for proofreading this manuscript.

## Author contributions

S.X. and T.H. conceived the research project. S.X. developed the computational algorithms for BUDDY. S.X., S.S., and B.X. constructed the standalone software platform in C#. S.X. performed method evaluations and applications. S.X. and T.H. wrote the manuscript, and all authors approved the final version.

---

**Extended Data Fig. 1.**
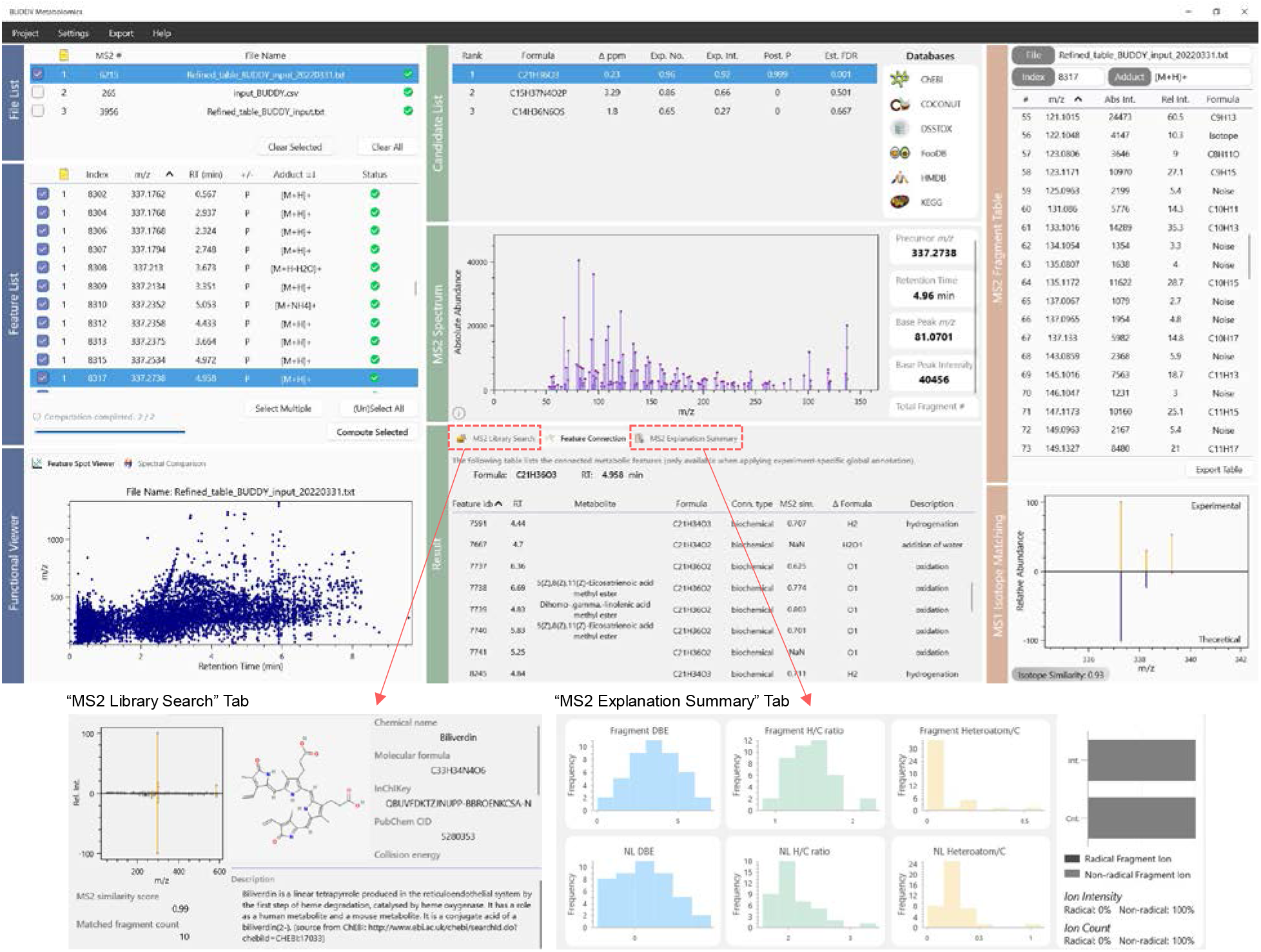
BUDDY graphical user interface. BUDDY offers an intuitive graphical user interface directly downloadable from GitHub. BUDDY can simultaneously process data files from different sources. For identified metabolites, MS/MS matching information and detailed chemical descriptions are provided to help check the identification quality and further investigate biological insights. If experiment-specific global peak annotation is performed, feature connections are listed with MS/MS similarity, formula difference, and connection description information.

**Extended Data Fig. 2.**
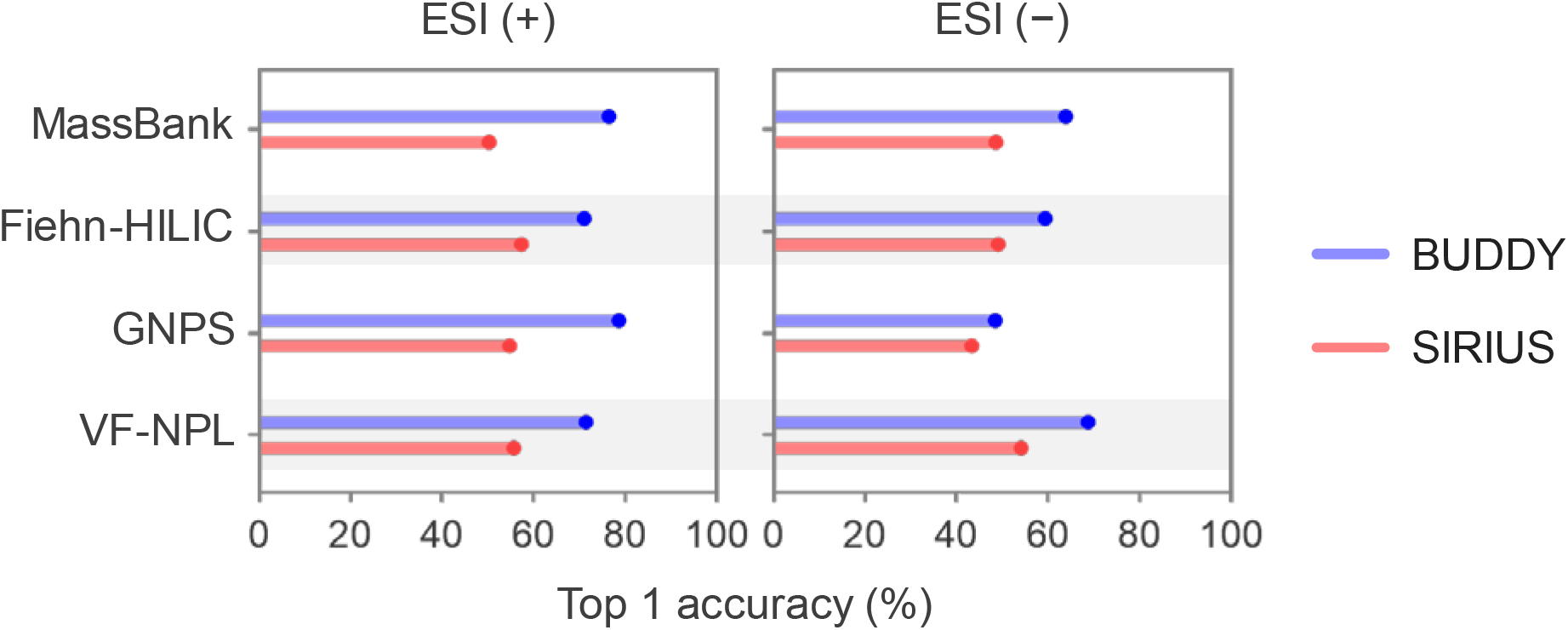
Method evaluation on MS/MS reference libraries (QTOF MS/MS). Top 1 accuracy of molecular formula annotation on four reference MS/MS libraries (QTOF MS/MS spectra shown). MS/MS and precursor *m/z* were used for annotation.

**Extended Data Fig. 3.**
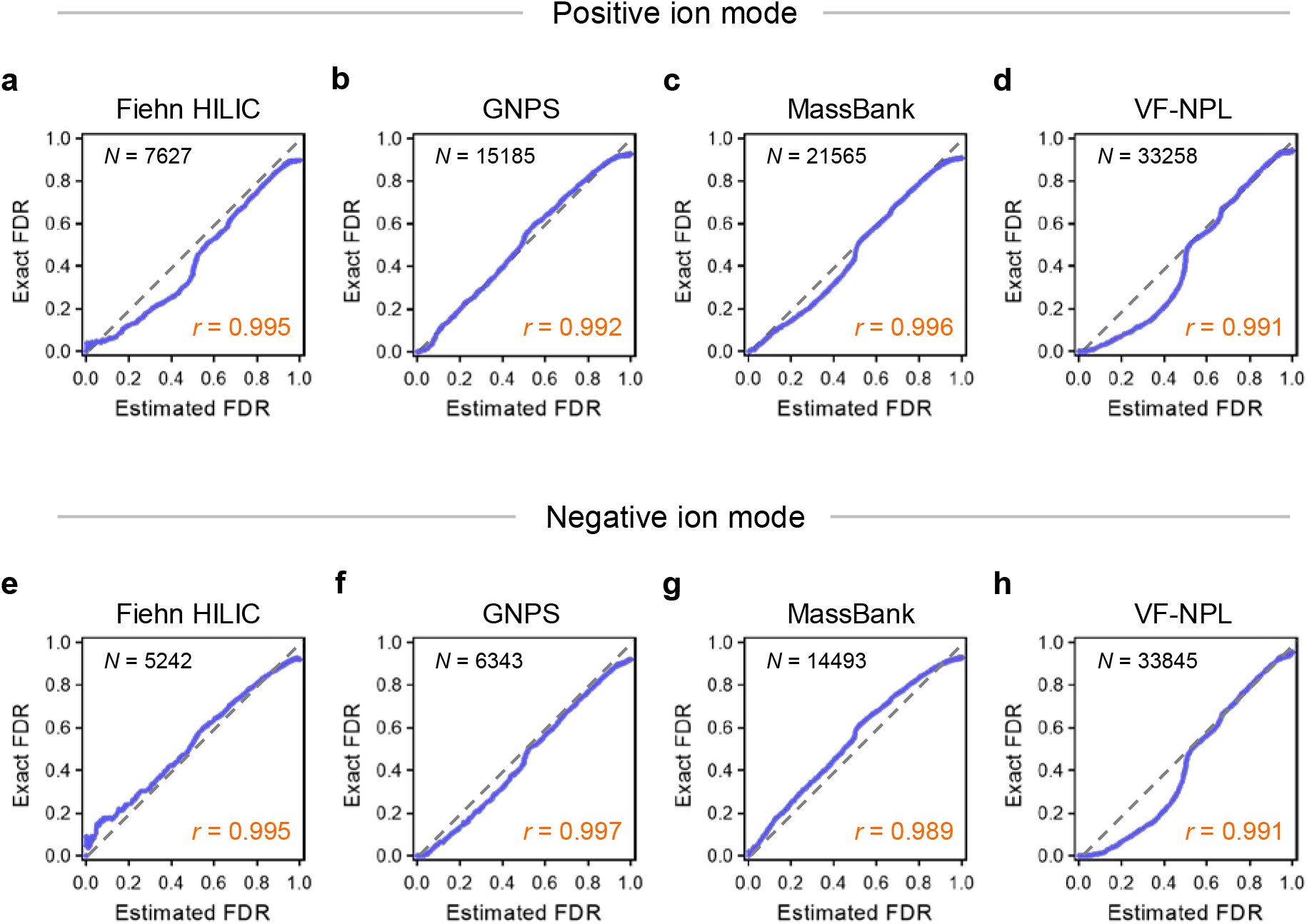
Validation of FDR estimation on reference MS/MS libraries. We used four reference MS/MS libraries to evaluate FDR estimation in BUDDY. Q-Q plots of estimated FDR and exact FDR are shown. In both positive and negative ion modes, estimated FDR shows high Pearson’s correlation coefficients (*r* >0.98) with exact FDR.

**Extended Data Fig. 4.**
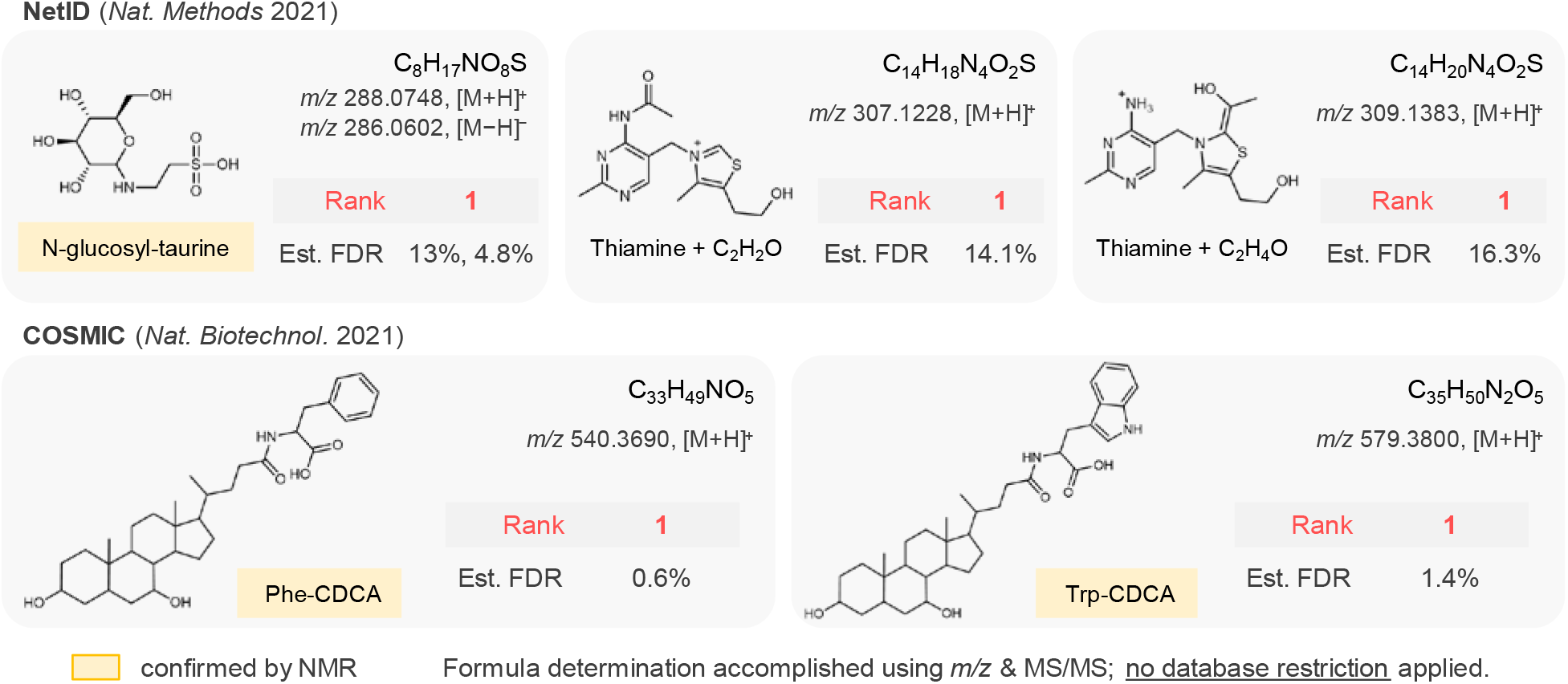
Method evaluation on novel compounds discovered in references. Bottom-up MS/MS interrogation was evaluated using the MS/MS spectra of five novel compounds, three of which were confirmed in NMR experiments. The correct formulae were ranked first in all cases, with the estimated FDR ranging from 0.6 to 16.3%.

**Extended Data Fig. 5.**
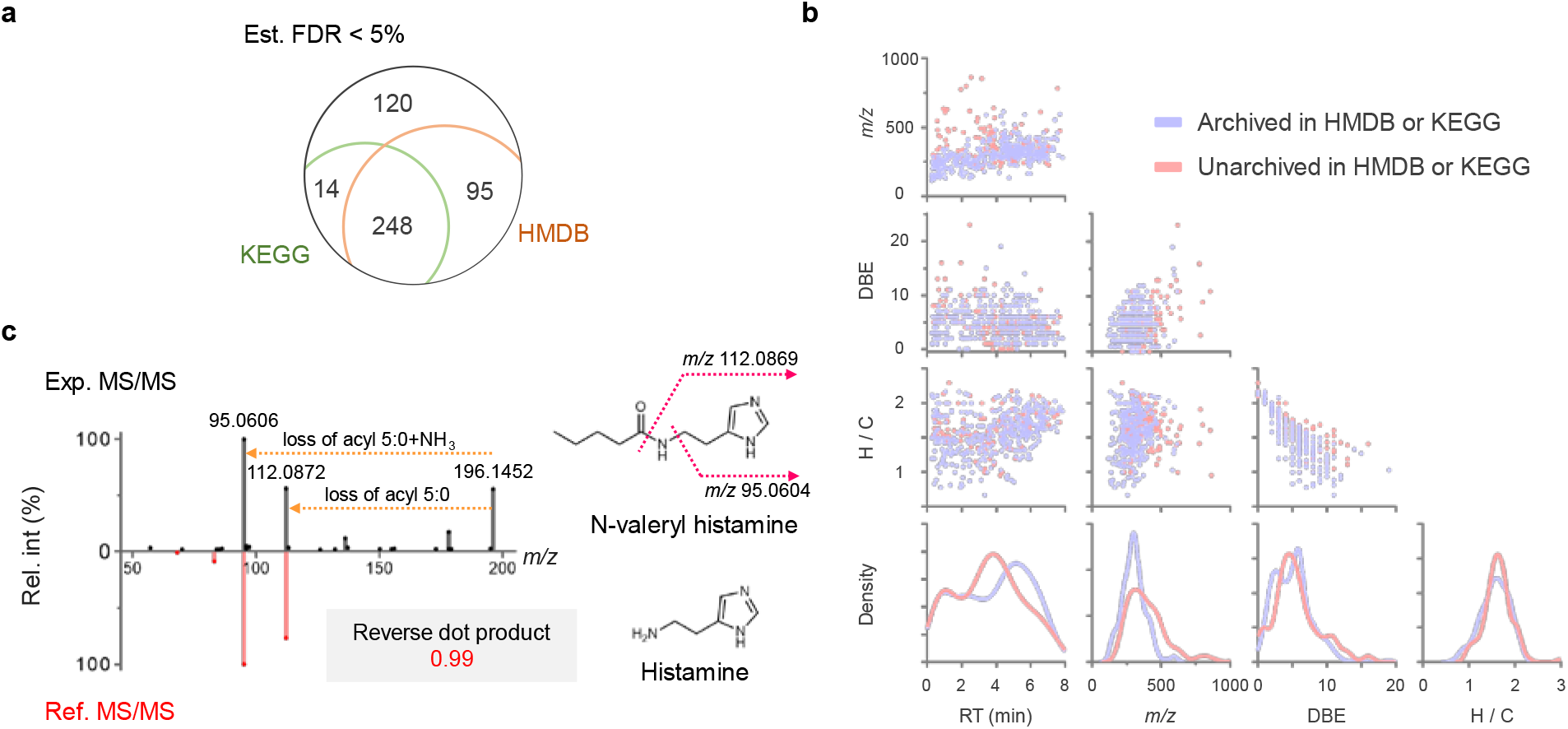
Molecular formula discovery in NIST human fecal material standards. **a**, Venn diagram of annotated molecular formulae with estimated FDR <5%. **b**, Scatter and density plots of *m/z,* retention time, DBE value, and hydrogen carbon ratio (H / C) of high-confidence annotated molecular formulae. **c**, Manual inspection of N-valeryl histamine. The experimental MS/MS of N-valeryl histamine shows a reverse dot product score of 0.99 against the reference MS/MS of histamine. Two neutral losses representing the acyl chain were found. Valeric acid is commonly produced in the gut microbiota (e.g., by Clostridia species), and histamine is a well-known diet-derived metabolite.

## References

1. Wang, M. et al. Sharing and community curation of mass spectrometry data with Global Natural Products Social Molecular Networking. Nature Biotechnology 34, 828–837 (2016).

2. NIST (2014).

3. Xue, J., Guijas, C., Benton, H.P., Warth, B. & Siuzdak, G. METLIN MS2 molecular standards database: a broad chemical and biological resource. Nature Methods 17, 953–954 (2020).

4. Horai, H. et al. MassBank: a public repository for sharing mass spectral data for life sciences. Journal of Mass Spectrometry 45, 703–714 (2010).

5. da Silva, R.R., Dorrestein, P.C. & Quinn, R.A. Illuminating the dark matter in metabolomics. Proceedings of the National Academy of Sciences 112, 12549 (2015).

6. Stein, S. Mass Spectral Reference Libraries: An Ever-Expanding Resource for Chemical Identification. Analytical Chemistry 84, 7274–7282 (2012).

7. Dührkop, K. et al. SIRIUS 4: a rapid tool for turning tandem mass spectra into metabolite structure information. Nature Methods 16, 299–302 (2019).

8. Bittremieux, W., May, D.H., Bilmes, J. & Noble, W.S. A learned embedding for efficient joint analysis of millions of mass spectra. Nature Methods 19, 675–678 (2022).

9. Dührkop, K., Shen, H., Meusel, M., Rousu, J. & Böcker, S. Searching molecular structure databases with tandem mass spectra using CSI:FingerID. Proceedings of the National Academy of Sciences 112, 12580 (2015).

10. Hoffmann, M.A. et al. High-confidence structural annotation of metabolites absent from spectral libraries. Nature Biotechnology (2021).

11. Chen, L. et al. Metabolite discovery through global annotation of untargeted metabolomics data. Nature Methods (2021).

12. Shen, X. et al. Metabolic reaction network-based recursive metabolite annotation for untargeted metabolomics. Nat Commun 10, 1516 (2019).

13. Ludwig, M. et al. Database-independent molecular formula annotation using Gibbs sampling through ZODIAC. Nature Machine Intelligence 2, 629–641 (2020).

14. Ernst, M. et al. MolNetEnhancer: Enhanced Molecular Networks by Integrating Metabolome Mining and Annotation Tools. Metabolites 9(2019).

15. Wishart, D.S. et al. HMDB 4.0: the human metabolome database for 2018. Nucleic Acids Research 46, D608–D617 (2018).

16. Kanehisa, M., Sato, Y., Kawashima, M., Furumichi, M. & Tanabe, M. KEGG as a reference resource for gene and protein annotation. Nucleic Acids Research 44, D457–D462 (2016).

17. Hastings, J. et al. ChEBI in 2016: Improved services and an expanding collection of metabolites. Nucleic acids research 44, D1214–1219 (2016).

18. Kim, S. et al. PubChem 2019 update: improved access to chemical data. Nucleic Acids Research 47, D1102–D1109 (2019).

19. Pence, H.E. & Williams, A. ChemSpider: An Online Chemical Information Resource. Journal of Chemical Education 87, 1123–1124 (2010).

20. Bocker, S. & Liptak, Z. A Fast and Simple Algorithm for the Money Changing Problem. Algorithmica 48, 413–432 (2007).

21. Böcker, S., Letzel, M.C., Lipták, Z. & Pervukhin, A. SIRIUS: decomposing isotope patterns for metabolite identification†. Bioinformatics 25, 218–224 (2009).

22. Rasche, F., Svatoš, A., Maddula, R.K., Böttcher, C. & Böcker, S. Computing Fragmentation Trees from Tandem Mass Spectrometry Data. Analytical Chemistry 83, 1243–1251 (2011).

23. Staden, R. A strategy of DNA sequencing employing computer programs. Nucleic Acids Research 6, 2601–2610 (1979).

24. Anderson, S. Shotgun DNA sequencing using cloned DNase I-generated fragments. Nucleic acids research 9, 3015–3027 (1981).

25. Aebersold, R. & Mann, M. Mass spectrometry-based proteomics. Nature 422, 198–207 (2003).

26. Chait, B.T. Mass spectrometry: bottom-up or top-down? Science (2006).

27. Scheubert, K. et al. Significance estimation for large scale metabolomics annotations by spectral matching. Nat Commun 8, 1494 (2017).

28. Pluskal, T., Uehara, T. & Yanagida, M. Highly Accurate Chemical Formula Prediction Tool Utilizing High-Resolution Mass Spectra, MS/MS Fragmentation, Heuristic Rules, and Isotope Pattern Matching. Analytical Chemistry 84, 4396–4403 (2012).

29. Xing, S. & Huan, T. Radical fragment ions in collision-induced dissociation-based tandem mass spectrometry. Analytica Chimica Acta 1200, 339613 (2022).

30. Senior, J.K. Partitions and Their Representative Graphs. American Journal of Mathematics 73, 663–689 (1951).

31. Platt, J. Probabilistic Outputs for Support Vector Machines and Comparisons to Regularized Likelihood Methods. Adv. Large Margin Classif. 10(2000).

32. Nikolskiy, I., Mahieu, N.G., Chen, Y., Jr., Tautenhahn, R. & Patti, G.J. An Untargeted Metabolomic Workflow to Improve Structural Characterization of Metabolites. Analytical Chemistry 85, 7713–7719 (2013).

33. Xing, S. et al. Recognizing Contamination Fragment Ions in Liquid Chromatography–Tandem Mass Spectrometry Data. Journal of the American Society for Mass Spectrometry 32, 2296–2305 (2021).

34. Djoumbou Feunang, Y. et al. ClassyFire: automated chemical classification with a comprehensive, computable taxonomy. Journal of Cheminformatics 8, 61 (2016).

35. Li, Y. et al. Spectral entropy outperforms MS/MS dot product similarity for small-molecule compound identification. Nature Methods 18, 1524–1531 (2021).

36. Schymanski, E.L. et al. Identifying small molecules via high resolution mass spectrometry: communicating confidence. Environ Sci Technol 48, 2097–2098 (2014).

37. Lai, Z. et al. Identifying metabolites by integrating metabolome databases with mass spectrometry cheminformatics. Nature Methods 15, 53–56 (2018).

38. Simón-Manso, Y. et al. Mass Spectrometry Fingerprints of Small-Molecule Metabolites in Biofluids: Building a Spectral Library of Recurrent Spectra for Urine Analysis. Analytical Chemistry 91, 12021–12029 (2019).

39. Wang, M. et al. Mass spectrometry searches using MASST. Nature Biotechnology 38, 23–26 (2020).

40. Cohen, L.J. et al. Commensal bacteria make GPCR ligands that mimic human signalling molecules. Nature 549, 48–53 (2017).

41. Chang, F.-Y. et al. Gut-inhabiting Clostridia build human GPCR ligands by conjugating neurotransmitters with diet- and human-derived fatty acids. Nature Microbiology 6, 792–805 (2021).

42. Giné, R. et al. HERMES: a molecular-formula-oriented method to target the metabolome. Nature Methods 18, 1370–1376 (2021).

43. Yin, Y., Wang, R., Cai, Y., Wang, Z. & Zhu, Z.-J. DecoMetDIA: Deconvolution of Multiplexed MS/MS Spectra for Metabolite Identification in SWATH-MS-Based Untargeted Metabolomics. Analytical Chemistry 91, 11897–11904 (2019).

44. Tsugawa, H. et al. MS-DIAL: data-independent MS/MS deconvolution for comprehensive metabolome analysis. Nature Methods 12, 523–526 (2015).

45. Tada, I. et al. Correlation-Based Deconvolution (CorrDec) To Generate High-Quality MS2 Spectra from Data-Independent Acquisition in Multisample Studies. Analytical Chemistry 92, 11310–11317 (2020).

46. Li, D. et al. XY-Meta: A High-Efficiency Search Engine for Large-Scale Metabolome Annotation with Accurate FDR Estimation. Analytical Chemistry 92, 5701–5707 (2020).

47. Bonini, P., Kind, T., Tsugawa, H., Barupal, D.K. & Fiehn, O. Retip: Retention Time Prediction for Compound Annotation in Untargeted Metabolomics. Analytical Chemistry 92, 7515–7522 (2020).

48. Bach, E., Szedmak, S., Brouard, C., Böcker, S. & Rousu, J. Liquid-chromatography retention order prediction for metabolite identification. Bioinformatics 34, i875–i883 (2018).

49. Domingo-Almenara, X. et al. The METLIN small molecule dataset for machine learning-based retention time prediction. Nature Communications 10, 5811 (2019).

50. Zhou, Z. et al. Ion mobility collision cross-section atlas for known and unknown metabolite annotation in untargeted metabolomics. Nature Communications 11, 4334 (2020).

51. Huber, F. et al. Spec2Vec: Improved mass spectral similarity scoring through learning of structural relationships. PLOS Computational Biology 17, e1008724 (2021).

52. Xing, S. et al. Retrieving and Utilizing Hypothetical Neutral Losses from Tandem Mass Spectra for Spectral Similarity Analysis and Unknown Metabolite Annotation. Analytical Chemistry 92, 14476–14483 (2020).

53. Treen, D.G.C. et al. SIMILE enables alignment of tandem mass spectra with statistical significance. Nature Communications 13, 2510 (2022).

54. van der Hooft, J.J.J., Wandy, J., Barrett, M.P., Burgess, K.E.V. & Rogers, S. Topic modeling for untargeted substructure exploration in metabolomics. Proceedings of the National Academy of Sciences 113, 13738 (2016).

55. Djoumbou-Feunang, Y. et al. BioTransformer: a comprehensive computational tool for small molecule metabolism prediction and metabolite identification. Journal of Cheminformatics 11, 2 (2019).

56. Jeffryes, J.G. et al. MINEs: open access databases of computationally predicted enzyme promiscuity products for untargeted metabolomics. Journal of Cheminformatics 7, 44 (2015).

57. Kind, T. & Fiehn, O. Seven Golden Rules for heuristic filtering of molecular formulas obtained by accurate mass spectrometry. BMC Bioinformatics 8, 105 (2007).

58. Ludwig, M. et al. Studying Charge Migration Fragmentation of Sodiated Precursor Ions in Collision-Induced Dissociation at the Library Scale. Journal of the American Society for Mass Spectrometry 32, 180–186 (2021).

59. Bertz, S.H. The first general index of molecular complexity. Journal of the American Chemical Society 103, 3599–3601 (1981).

60. Ertl, P., Roggo, S. & Schuffenhauer, A. Natural Product-likeness Score and Its Application for Prioritization of Compound Libraries. Journal of Chemical Information and Modeling 48, 68–74 (2008).

61. Kessner, D., Chambers, M., Burke, R., Agus, D. & Mallick, P. ProteoWizard: open source software for rapid proteomics tools development. Bioinformatics 24, 2534–2536 (2008).

62. Stein, S.E. & Scott, D.R. Optimization and testing of mass spectral library search algorithms for compound identification. Journal of the American Society for Mass Spectrometry 5, 859–866 (1994).

