## Supplementary information for "Molecular formula discovery via bottom-up MS/MS interrogation"

Website: <https://huan.chem.ubc.ca/>

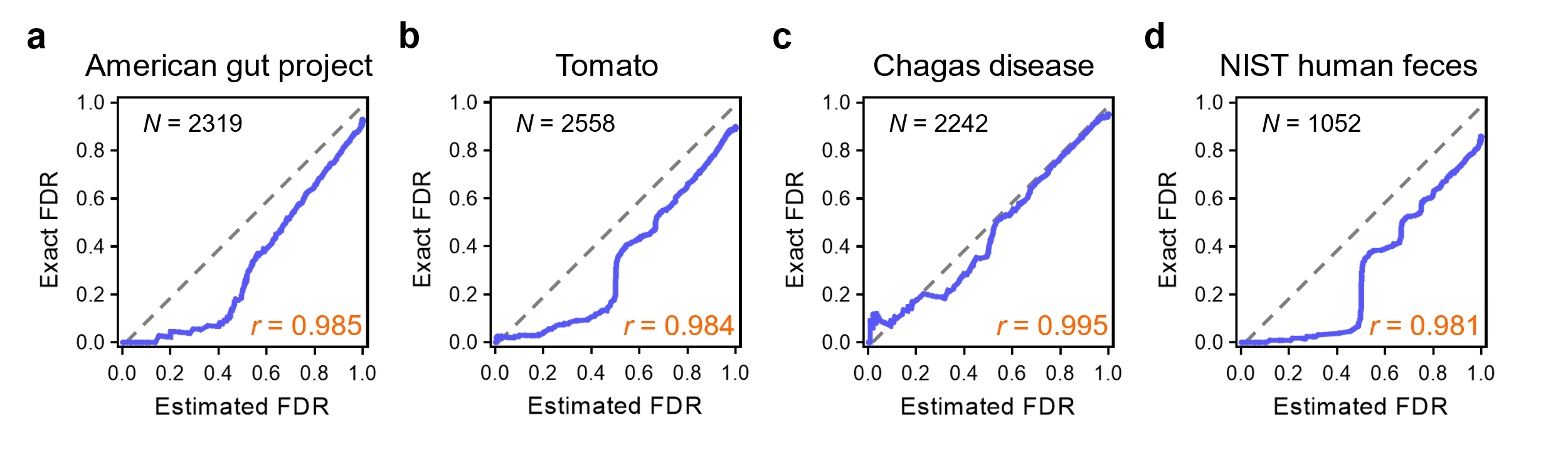

### Supplementary Fig. 1 | FDR estimation on LC-MS datasets. We used four publicly available LC-MS datasets to evaluate FDR estimation in BUDDY. Q-Q plots of estimated FDR and exact FDR are shown. In all tested LC-MS datasets, estimated FDR shows Pearson’s correlation coefficients of >0.98 with exact FDR.

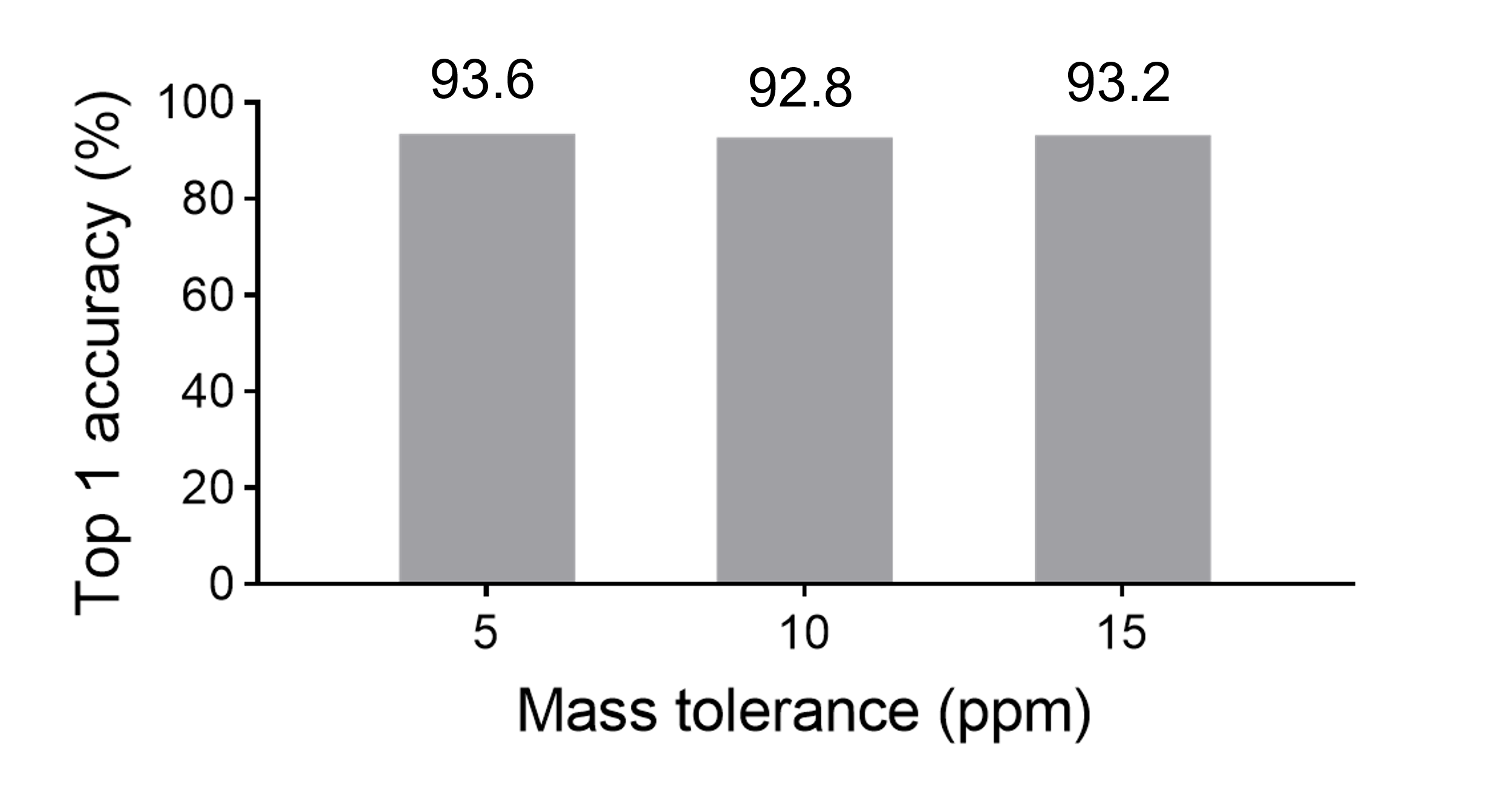

### Supplementary Fig. 2 | Experiment-specific mass accuracy estimation. Shown in the column plot are the top 1 accuracies of molecular formula annotation on the tomato dataset using different mass tolerances. A common acceptable mass tolerance is 5 ppm for Orbitrap MS instruments, but annotation accuracy only drops by 0.4% to 93.2% even when 15 ppm is used.

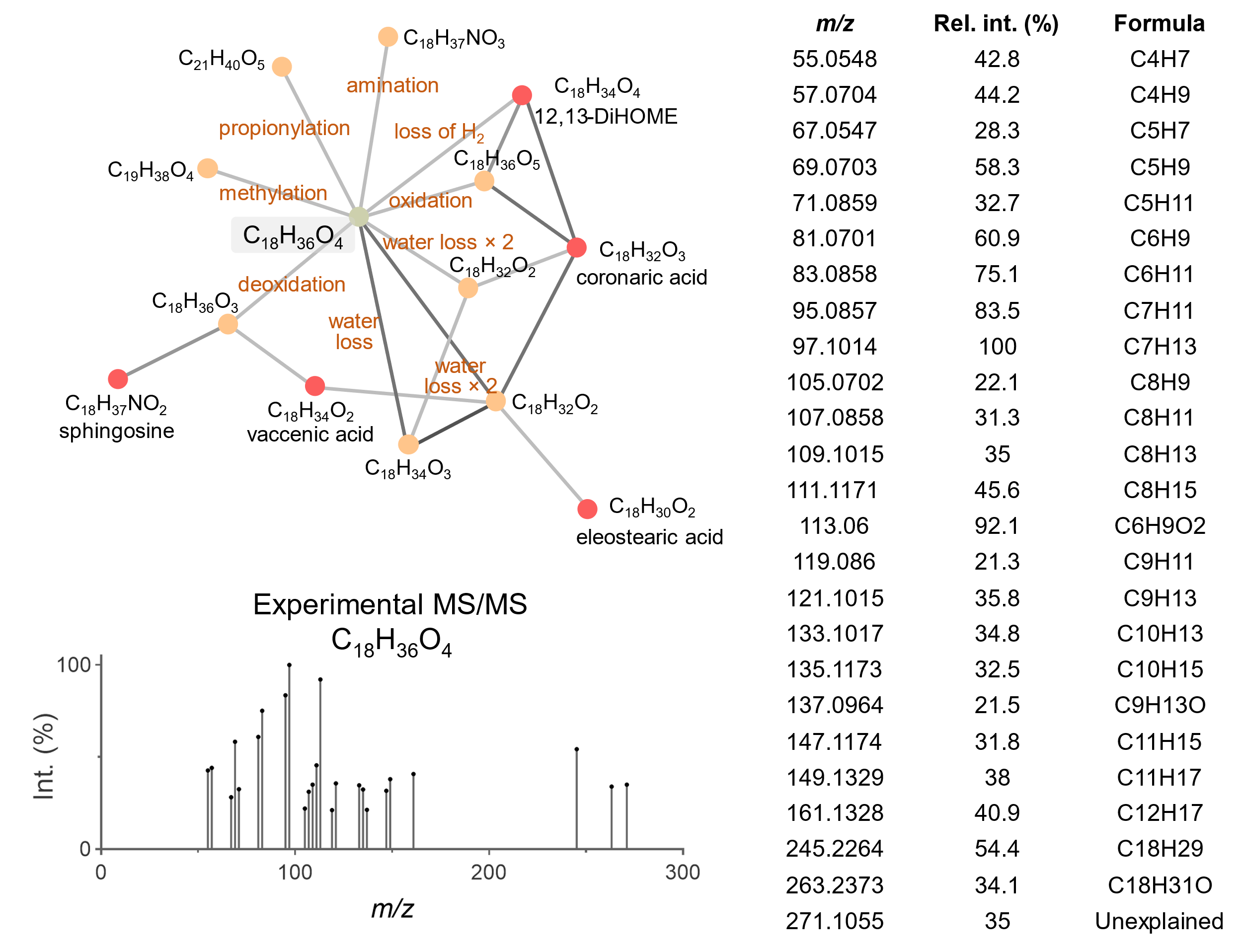

### Supplementary Fig. 3 | Subformula annotations of target feature MS/MS fragments. The surrounding molecular network indicates that the structure of the target feature is a dihydroxylated long-chain fatty acid. Subformula annotations of its MS/MS fragments also imply the existence of a long acyl group.

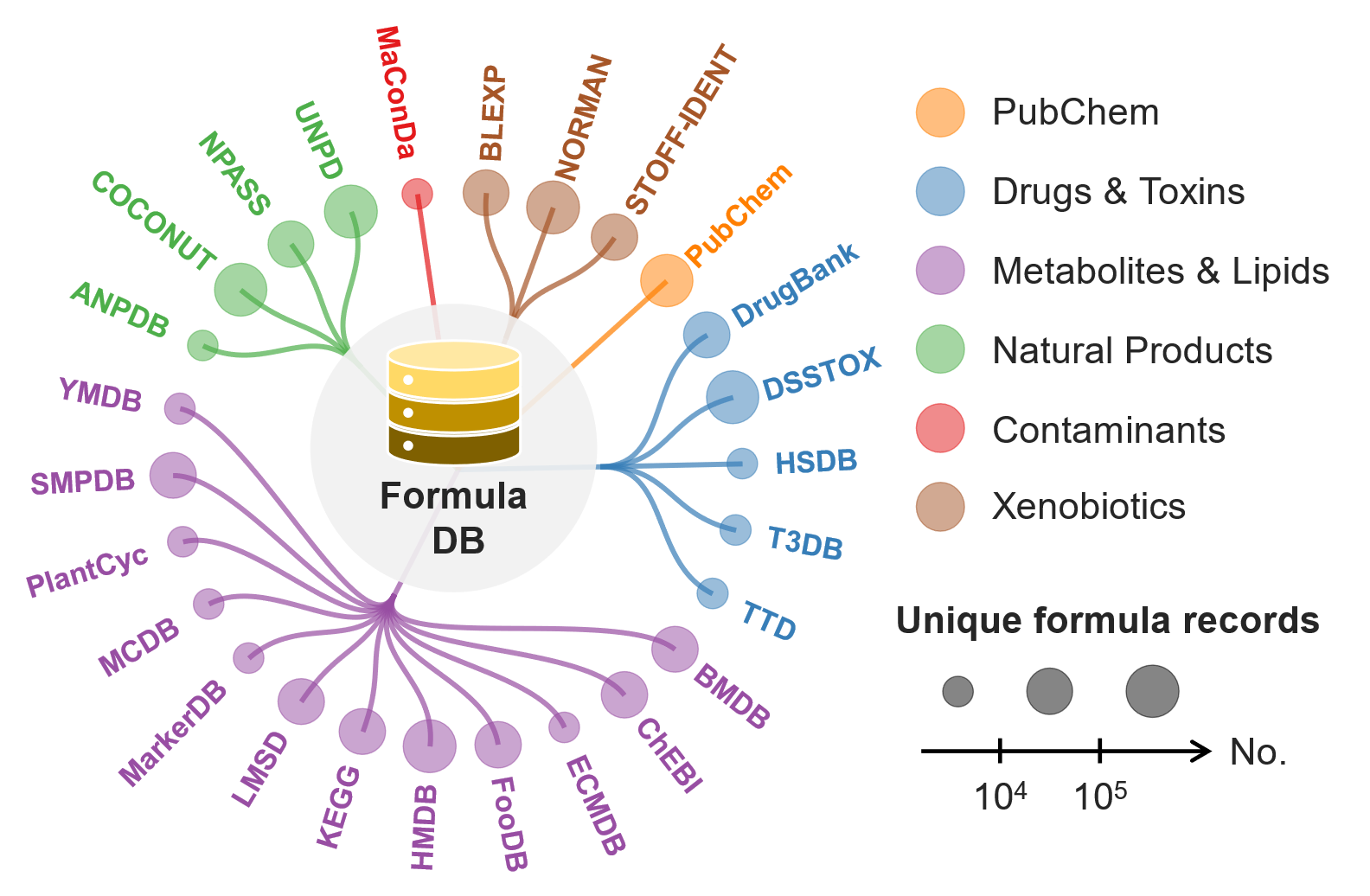

### Supplementary Fig. 4 | The curated formula database archiving >3.5 million unique molecular formulae from 26 chemical databases. The 26 chemical databases representing drugs & toxins, metabolites & lipids, natural products, contaminants, and xenobiotics were integrated and curated. The entire chemical compound repository in PubChem was also included.

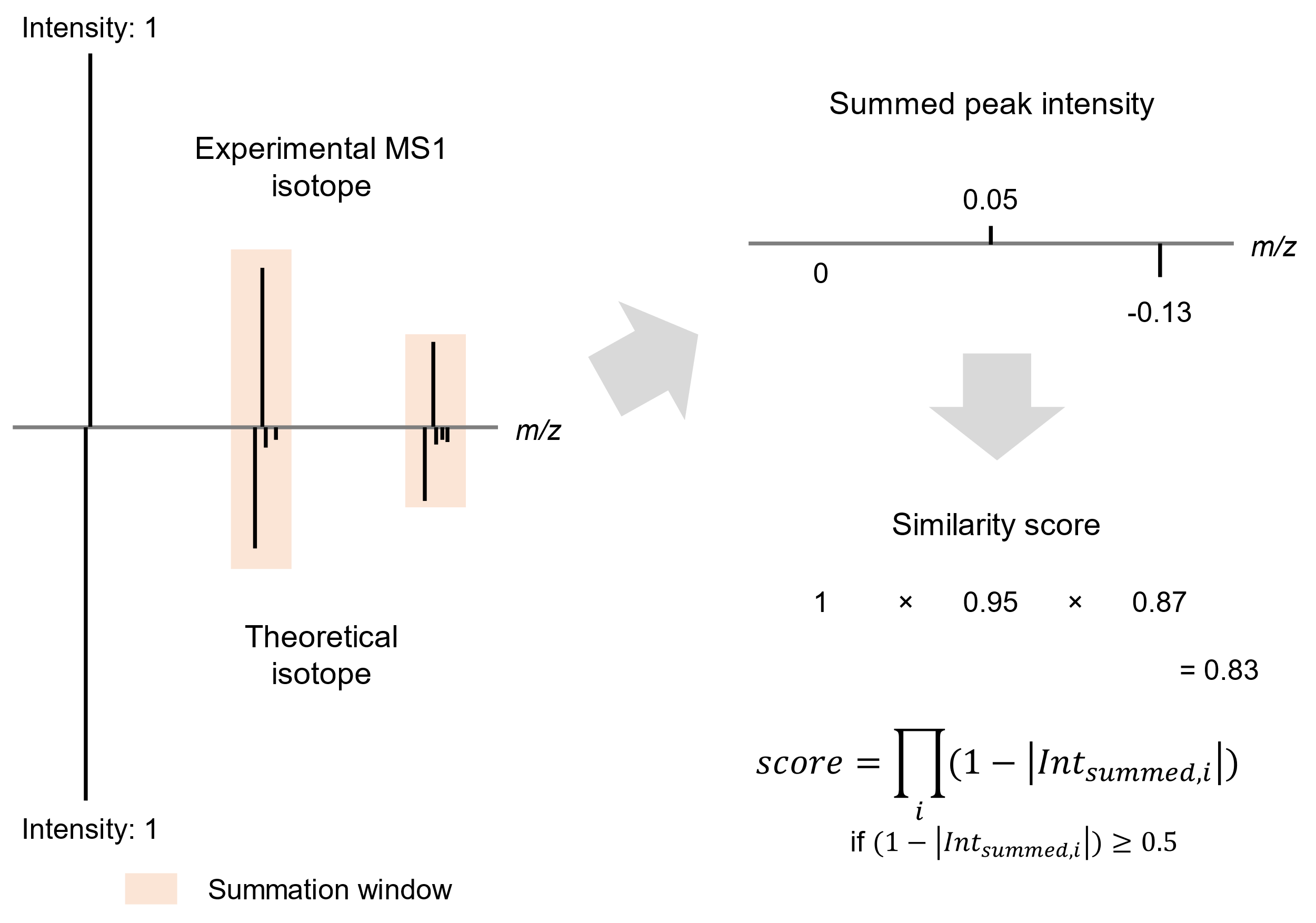

### Supplementary Fig. 5 | Illustration of the isotope similarity algorithm applied in BUDDY. Intensity-normalized isotopic patterns are subjected to peak intensity summation and score calculation. The final isotope similarity score is calculated as the product of each pair of summed peak intensities. Notably, we set the threshold of $\boldsymbol{1-}\left| \boldsymbol{Int}_{\boldsymbol{summed,i}} \right|$ as 0.5 to reduce the potential of including coeluted ions into the isotope similarity scoring, which will dramatically diminish the final score. This means if $\boldsymbol{1-}\left| \boldsymbol{Int}_{\boldsymbol{summed,i}} \right|\boldsymbol{<0.5}$ is true for peak pair *k*, the final score calculation stops, and no more peaks (*k*, *k*+1, *k*+2, etc.) will be considered.

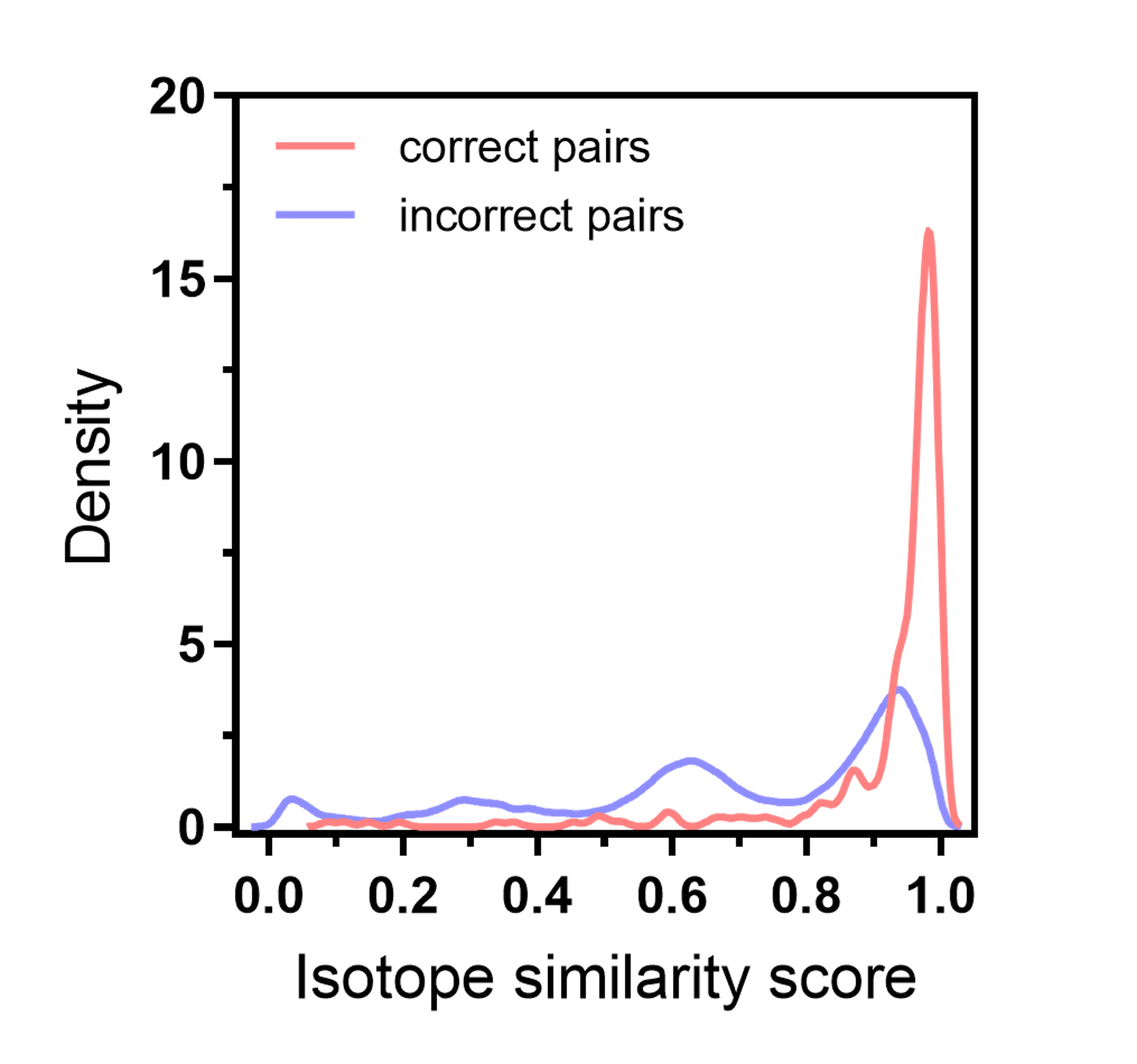

### Supplementary Fig. 6 | Validation of the isotope similarity algorithm. Both two-sample Kolmogorov–Smirnov test (two-sided, *P* < 2.2×10^−16^) and Mann-Whitney U test (two-sided, *P* < 2.1×10^−103^) show statistical significances between the isotope similarity scores of correct pairs and incorrect pairs.

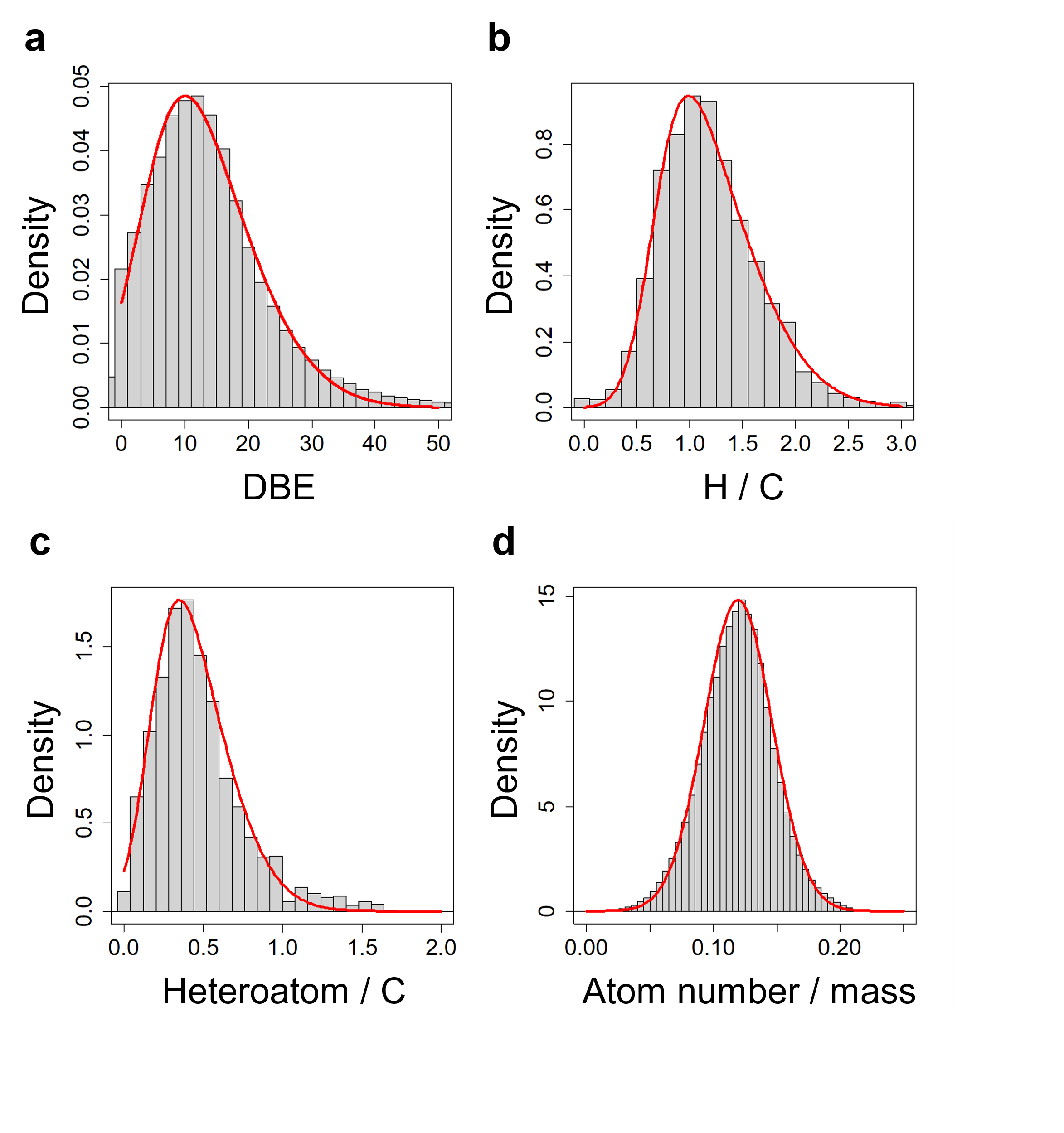

### Supplementary Fig. 7 | Neutral formula database distribution analysis. a, The distribution of DBE values. b, The distribution of hydrogen / carbon ratios. c, The distribution of heteroatom / carbon ratios. d, The distribution of atom number / mass ratios.

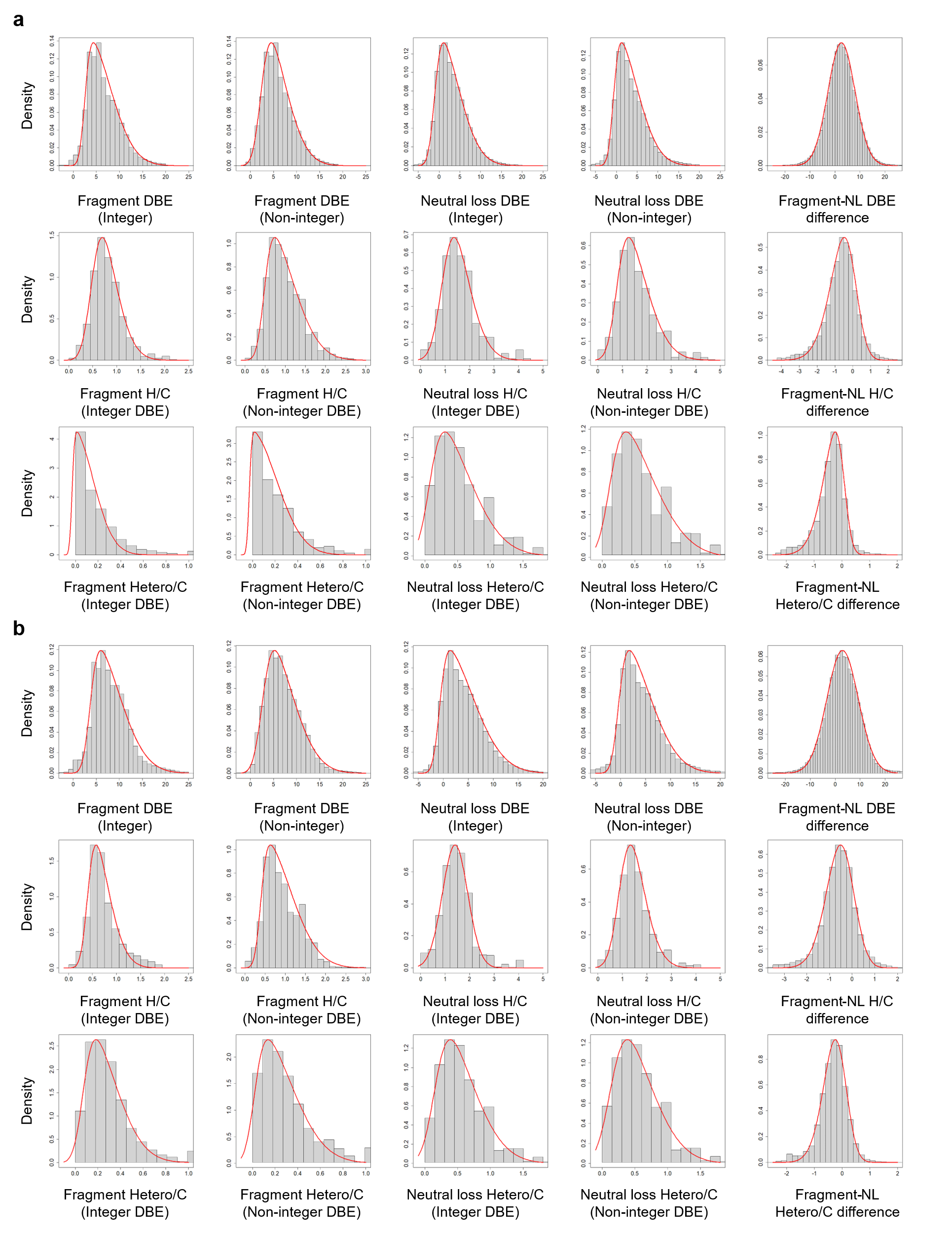

### Supplementary Fig. 8 | NIST20 distribution analysis. A total of 15 MLR features’ distributions were analyzed, plotted, and fitted with the skew normal distribution. a, NIST20 positive ion mode. b, NIST20 negative ion mode.

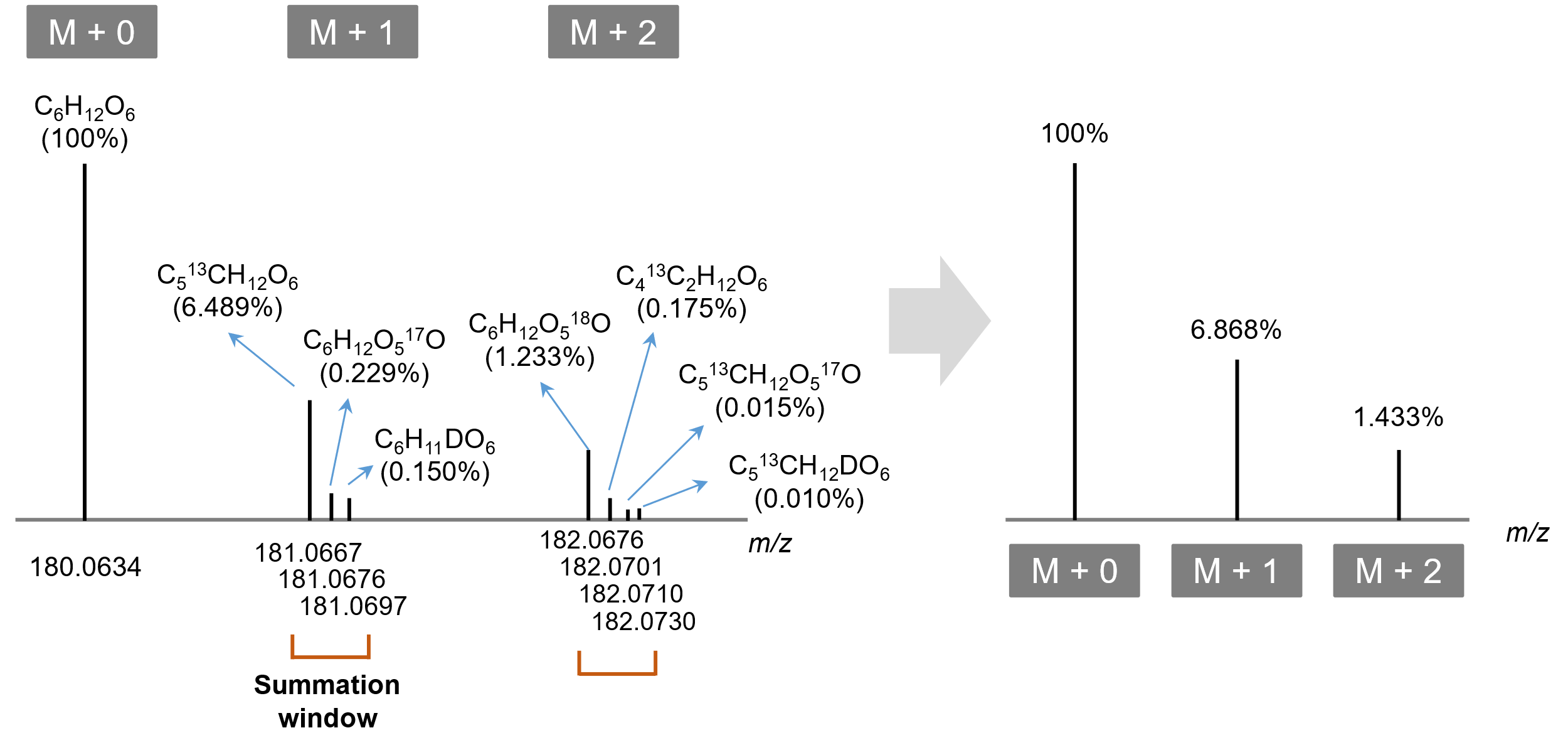

### Supplementary Fig. 9 | Illustration of isotopologue grouping in the isotope simulation process. The isotope intensities for adjacent isotopologues were summed within a given summation window. Details are discussed in Supplementary Note 9.

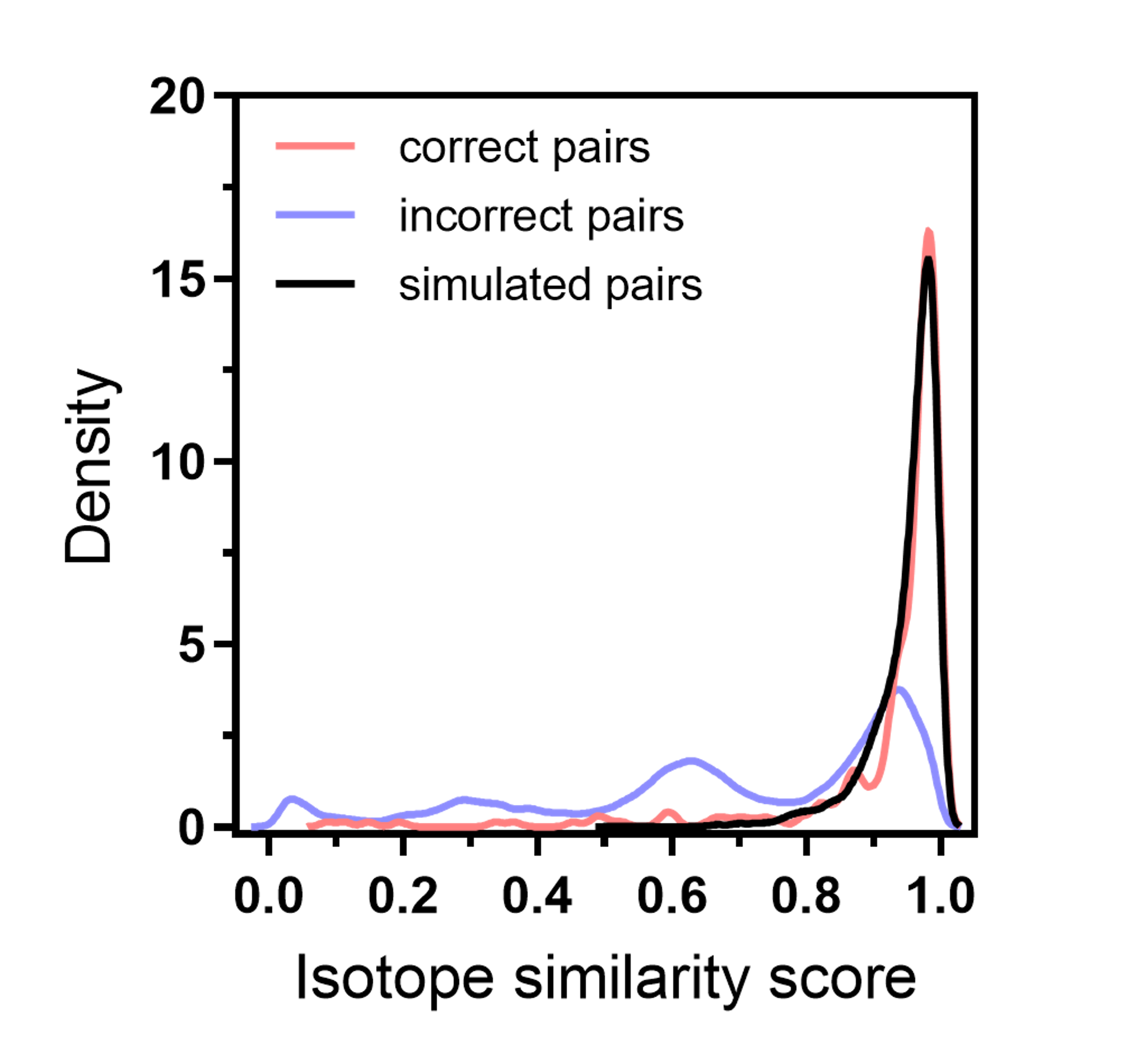

### Supplementary Fig. 10 | Validation of the isotope simulation workflow. No statistical significance can be observed between the isotope similarity score distributions of “simulated pairs” and “correct pairs” (two-sided two-sample Kolmogorov–Smirnov test: *P* > 0.10; two-sided Mann-Whitney *U* test: *P* > 0.70).

### Supplementary Note 1 | Manual inspection of a newly discovered molecular formula.

The manual inspection of a novel formula (*m/z* 674.4387, C_35_H_64_NO_9_P) is illustrated in **Fig. 5b**. In its MS/MS spectrum, we found the characteristic fragment ion (*m/z* 184.0733, C_5_H_15_NO_4_P^+^) representing glycerophosphocholine (PC) lipids. Two side chains were confirmed via their corresponding neutral losses, one of which is an oxidized fatty acyl chain (9-oxo-nonanoic acid). As described in HMDB, 9-oxo-nonanoic acid is a human metabolite that can be synthesized into PC(16:0/9:0(CHO)). Oxidized PCs, however, are neither broadly discovered nor archived in current chemical databases; the LIPID MAPS Structure Database^1^, a comprehensive database for lipids, only contains two 9-oxononanoyl PCs: (PC(16:0/9:0(CHO)) and PC(18:1/9:0(CHO))). With the aid of BUDDY, we are the first to report PC(18:2/9:0(CHO)), implying the validity of novel formulae annotated at low FDR.

### Supplementary Note 2 | Orthogonal evaluation of ARUS MS/MS libraries.

The method performance was verified in an orthogonal manner. ARUS MS/MS libraries provide spectral comparison results using the hybrid similarity search^2^ (HSS) to search for structural analogs within the MS/MS reference library (NIST17 in this case). In cases where precursor mass differences match certain biochemical transformations, formula annotation for unknown MS/MS can be achieved by biochemical modifications on known chemical standards. To illustrate (**Fig. 5c**), an unidentified spectrum with precursor *m/z* 448.3063 has a matching score of 993 (out of 999) against the reference spectrum of 12-ketodeoxycholic acid (molecular formula: C_24_H_38_O_4_, *m/z* 391.2843), and their precursor mass difference (57.0220 Da) can be identified as glycine conjugation (C_2_H_3_NO). The formula of the unidentified MS/MS can thus be inferred as C_26_H_41_NO_5_ (C_24_H_38_O_4_ + C_2_H_3_NO). This result is in accord with BUDDY’s annotation (estimated FDR: 1.3%). Searching this MS/MS against the entire public repository using MASST^3^, the results suggest that this bile acid derivative is commonly detected in 28 metabolomic datasets collected from humans, mice, and bacteria. Also, glycine-conjugated bile acids have been reported in humans and avian species in recent decades^4^, further supporting its structural existence. However, it is noteworthy that not even its molecular formula is archived in HMDB or KEGG, implying the urgent necessity of exploring novel formulae beyond current metabolite databases. As an example of xenobiotic compounds detected in humans, BUDDY agreed with the spectral matching approach on hydroxy-zolpidem, which is also absent from both HMDB and KEGG (**Fig. 5d**). A clear mass shift of oxygen can be observed comparing the unknown MS/MS with the reference spectrum of zolpidem. Zolpidem is known as a hypnotic drug designed for patients with insomnia, and revealing zolpidem-related metabolites is essential for drug metabolism studies and forensic tests^5^. BUDDY correctly assigned its molecular formula at a low estimated FDR of 1.8% without the assistance of MS/MS reference libraries. Finally, as a large-scale orthogonal test, we focused on the ARUS MS/MS with HSS spectral matching scores of ≥950. We reserved ARUS MS/MS of precursors that have mass differences with reference spectra matching biochemical transformations. Formula determination was completed using spectral matching and BUDDY in parallel. In both approaches, 90.3% (436 out of 483 tested MS/MS spectra) resulted in the same formula (**Supplementary Table 10**). Further considering the MS/MS spectra annotated with high confidence (<5% FDR) in BUDDY, 97.9% (284 out of 290) of tested spectra yielded consistent results. Notably, the above results were generated without MS1 isotope information, and a better performance can be anticipated in real metabolomic applications.

### Supplementary Note 3 | Experiment-specific mass deviation estimation.

BUDDY is able to estimate experiment-specific mass deviations to achieve more accurate predictions. This function automatically activates during the experiment-specific global peak annotation.

Metabolomics practitioners usually set up an empirical mass tolerance based on the MS instrument type when carrying out downstream data analyses. However, practical mass accuracy varies from experiment to experiment considering the mass spectral resolving power at different mass ranges and the ion signal level. To illustrate, different sample types may have different mass ranges (metabolites versus lipids), and larger masses usually have larger relative mass deviations, which leads to different mass deviation distributions among experiments.

We thus designed a workflow for experiment-specific mass deviation estimation such that the data input of different experiments can be standardized (normalized), which benefits the formula prediction task. In other words, the distribution of observed mass deviations varies from experiment to experiment, and it could stray from expectations while training the MLR models. This difference would diminish the prediction performance of our trained MLR models on specific experiment datasets. As such, we apply experiment-specific MLR feature scaling prior to MLR predictions. Details are as shown below.

We first begin with how mass deviation is usually defined and how it is implemented in BUDDY. Given the MS1 mass tolerance ${mass}_{tolerance}$ (in ppm), the mass of detected precursor ${mass}_{precursor}$, and the theoretical mass of ground truth formula ${mass}_{ground truth}$, the relative mass deviation $\Delta mass (ppm)$ is then:

$$\Delta mass (ppm)=\frac{{mass}_{precursor}-{mass}_{ground truth}}{{mass}_{ground truth}}\times{10}^{6}$$

During the MLR model training phase, we estimate that the mass deviation (in ppm) conforms to a Gaussian distribution with a mean of 0 and SD of 1/5 of the commonly used mass tolerance (MS instrument-specific, e.g., Orbitrap: 5 ppm), such that

$${\Delta mass}_{training}\sim N(0,{(\frac{{mass}_{tolerance}}{5})}^{2})$$

Note that $\Delta mass (ppm)$ is the mass deviation of the detected precursor mass relative to the theoretical mass given that the ground truth formula is known. In the case of searching for correct formulae given the observed precursor mass, $\Delta mass (ppm)$ should be calculated using ${mass}_{precursor}$ as a template. Thus, given the theoretical mass of a candidate formula ${mass}_{candidate}$, the mass deviation $\Delta mass (ppm)$ can be modified as

$$\Delta mass (ppm)=\frac{{mass}_{precursor}-{mass}_{candidate}}{{mass}_{precursor}}\times{10}^{6}$$

Additionally, $\Delta mass (ppm)$ can be positive or negative; we care more about its absolute value and how it compares to ${mass}_{tolerance} (ppm)$. We thus designed an MLR feature associated with precursor mass deviation, $precursorMzErrorRatio$. The $precursorMzErrorRatio$ for a candidate is calculated as

$$precursorMzErrorRatio=\frac{\left| \Delta mass \right|}{{mass}_{tolerance}}$$

Naturally, $precursorMzErrorRatio$ is a numeric value between 0 and 1, as all formula candidates are searched within the given MS1 mass tolerance range ($\left| \Delta mass \right|\leq{mass}_{tolerance}$).

Recall that in the training phase, the relative mass deviation conforms to a normal distribution, ${\Delta mass}_{training}\sim N(0,{(\frac{{mass}_{tolerance}}{5})}^{2})$, so the distribution of ${precursorMzErrorRatio}_{training}$ can be inferred as

$${precursorMzErrorRatio}_{training}=\frac{\left| {\Delta mass}_{training} \right|}{{mass}_{tolerance}}\sim\left| N(0,{(\frac{1}{5})}^{2}) \right|$$

In the experimental data, ${\Delta mass}_{experimental}$ for correct annotations conforms to another normal distribution with a SD of $\alpha\times{mass}_{tolerance}$

$${\Delta mass}_{exp, correct}\sim N(0,{(\alpha\times{mass}_{tolerance})}^{2})$$

where $\alpha$ is experiment-specific. Similarly, we have

$${precursorMzErrorRatio}_{exp, correct}\sim\left| N(0,\alpha^{2}) \right|$$

For incorrect annotations, ${\Delta mass}_{exp,incorrect}$ and ${precursorMzErrorRatio}_{exp,incorrect}$ can be modelled as uniform distributions.

$${precursorMzErrorRatio}_{exp, incorrect}\sim U(0,\beta)$$

Thus, the observed $precursorMzErrorRatio$ in an entire experimental dataset is the summation of two distributions. To correct the observed distribution back to the training condition, we defined the correction factor using the medians of the ${precursorMzErrorRatio}_{training}$ and the observed ${precursorMzErrorRatio}_{experimental}$.

$$Correction factor={[\frac{median({precursorMzErrorRatio}_{training})}{median({precursorMzErrorRatio}_{experimental})}]}^{\lambda}$$

Each $precursorMzErrorRatio$ value in this LC-MS run can be corrected as

$${precursorMzErrorRatio}_{corrected}={precursorMzErrorRatio}_{original}\times Correction factor$$

where $\lambda=0.5$, and $median\left( {precursorMzErrorRatio}_{training} \right)$ can be obtained during the training process.

Such a correction can better estimate the real distribution of experimental mass deviations and apply MLR models in a condition that is closer to the training condition.

### Supplementary Note 4 | Automated MS/MS noise peak elimination.

To ensure the spectral quality of the input experimental MS/MS spectra in BUDDY, we provide an optional step of automated MS/MS noise peak elimination. The detailed workflow is provided as follows.

Three user-defined parameters are $maxNoiseFragmentRatio$(default: 80%), $maxNoiseRSD$ (default: 0.25), and $maxNoiseIntensity$ (default: 5e3).

Given an MS/MS spectrum with *m* fragments, noise peak elimination only proceeds if $m>10$. MS/MS fragments are pre-sorted by peak intensity. The lowest 3 fragments are selected first, and the relative standard deviation (RSD) value of their peak intensities is calculated as $tmpRSD$. The temporary intensity threshold $tmpThreshold$ is calculated using the selected peaks as $Mean(intensity)+3\times SD(intensity)$. We then iteratively update the selected peak group. For the *i*-th loop ($1\leq i\leq20$), the selected peak group is updated as the lowest *k* fragments, where $k=roundToInteger(5\%\times i\times m)$.

Meanwhile, $tmpRSD$ and $tmpThreshold$ keep updating until one of the following three conditions is met:

a) $tmpRSD\geq maxNoiseRSD$ (RSD exceeds the defined threshold)

b) $tmpThreshold \geq maxNoiseIntensity$ (maximum noise peak intensity exceeds the defined threshold)

c) $5\%\times i\times m\geq maxNoiseFragmentRatio$ (maximum noise peak ratio exceeds the defined threshold)

The last updated $tmpThreshold$ value serves as the final peak intensity cutoff, and MS/MS peaks with intensities lower than the cutoff are discarded to complete the noise peak elimination process.

### Supplementary Note 5 | Automated MS/MS isotope peak recognition.

To remove the unwanted MS/MS isotope peaks (e.g., M + 1, M + 2), an optional step of automated MS/MS isotope peak recognition is provided.

Given a mass tolerance ${mass}_{tol}$ and fragment A $({m/z}_{A},{intensity}_{A})$, if there is another fragment B $({m/z}_{B},{intensity}_{B})$ where both $\left| {m/z}_{B}-({m/z}_{A}+1.003355) \right|\leq{mass}_{tol}$ and ${intensity}_{B}<{intensity}_{A}$ are satisfied, fragment B is identified as an isotope peak. All isotope peaks are first recognized and then discarded before further formula annotation. We herein use the mass difference between ^13^C (13.003355 Da) and ^12^C (12 Da) as the mass of a neutron, as carbon tends to contribute the most to M + 1 peaks. On the other hand, we use a relatively wide mass tolerance for isotope peak recognition considering the neutron mass difference among different elements (e.g., $mass\left( {}_{1}^{2}H \right)-mass\left( {}_{1}^{1}H \right)>mass\left( {}_{6}^{13}C \right)-mass({}_{6}^{12}C)$). We acknowledge this isotope peak recognition algorithm does not work perfectly for chlorine- or bromine-containing formulae as most of their isotopic peaks do not follow the conventional monotonically decreasing pattern. However, as an automated isotope recognition program, it is sufficient for removing isotope peaks in MS/MS spectra for most CHNOPS-containing formulae.

### Supplementary Note 6 | Radical fragment ions.

An important feature of BUDDY lies in its automatic recognition of radical fragment ions in MS/MS spectra. A recent study^6^ has demonstrated the unexpected frequency of radical fragment ions during the collision-induced dissociation (CID) process. From the aspect of algorithmic design, BUDDY is able to recognize radical (odd-electron) fragment ions at the same scale as even-electron fragment ions. When annotating fragment ions, two modified formula databases are searched simultaneously, an even-electron fragment ion formula database and a radical fragment ion formula database. On the basis of the neutral formula database, the formulae and masses in the even-electron fragment ion formula database are adjusted by a proton (M → [M + H]^+^ / [M – H]^–^); the formulae in the radical fragment ion formula database remain unchanged while the masses are adjusted by an electron (M → [M]^+•^ / [M]^–•^). As such, BUDDY shows no bias in recognizing radical fragment ions and even-electron fragment ions. Similar approaches are also applied to neutral loss searching. During the formula stitching process, even-electron fragment ions only pair with even-electron neutral losses; odd-electron fragment ions only pair with odd-electron neutral losses.

### Supplementary Note 7 | Evaluation of the MS1 isotope similarity algorithm.

We first collected the experimental MS1 isotopic patterns of 316 chemical standards (**Supplementary Table 16**). For each chemical standard, we first computed the isotope similarity score between its experimental isotope and the theoretical isotope; these isotope similarity scores are referred to as "correct pairs”. Then, we generated isotope similarity scores for “incorrect pairs”. For each chemical, we searched its mass against the formula database (mass searching tolerance: 10 ppm) and retrieved all molecular formulae other than its own to produce an “incorrect formula pool”. Isotope similarity scores of “incorrect pairs” were computed between experimental isotopes of chemical standards and theoretical isotopes of the “incorrect formula pool”. Between the isotope similarity score distributions of “correct pairs” and “incorrect pairs” (**Supplementary Fig. 6**), statistically significant differences were observed in both median values (two-sided Mann-Whitney *U* test, *P* < 3.7×10^−102^) and empirical distribution functions (two-sided two-sample Kolmogorov–Smirnov test, *P* < 2.2×10^−16^). Similar results could be obtained even using a narrower mass searching window (mass searching tolerance: 5 ppm; two-sided Mann-Whitney *U* test: *P* < 1.9×10^−101^; two-sided Kolmogorov–Smirnov test: *P* < 2.2×10^−16^), demonstrating the credibility of the isotope similarity calculation algorithm.

### Supplementary Note 8 | Distribution analyses guiding MLR feature generation.

***Neutral formula database distribution analysis.*** In order to guide the MS1-related feature generation in machine-learned ranking (MLR), we systematically analyzed the neutral formula database on a large scale. Inspired by ref. ^7^, we included the following candidate formula-relevant MLR features: DBE value, hydrogen/carbon ratio, heteroatom/carbon ratio, and total atom number/mass ratio. For the above four MLR features, we plotted their distributions using all valid neutral formulae in the database and fit the distribution curves individually with the skew normal distribution (**Supplementary Table 18 & Supplementary Fig. 7**). When mapping these MLR features onto the MLR feature matrix, we took the common logarithm of the two-sided area under the skewed Gaussian curve, log(*Area*). Our training data, however, cannot cover the entire possible ranges of DBE values and various ratios, meaning that there can be cases of experimental inputs exceeding the boundaries built during the model training process. As such, we set the lower limit of log(*Area*) as −2 to tolerate all possible extreme cases, which refers to a *P* value of 0.01.

***NIST20 distribution analysis.*** NIST20 was analyzed to guide the MS/MS-related feature generation in the MLR task. In short, we referred to the MS/MS spectra with fragment formulae annotated in NIST20 and constructed the distribution curves for pre-designated MS/MS-related features. Details about subformula annotation of fragment ions in NIST20 can be found in ref. ^8^. NIST20 was curated as follows. First, MS/MS spectra with common adduct forms were selected. We removed MS/MS spectra with isotopic adducts (e.g., MS/MS spectra for M + 1 ions), non-singly charged adducts, or adducts containing chemical elements beyond the allowed elemental list (e.g., [M + Li]^+^). Next, MS/MS spectra without InChIKey information or annotated subformulae were removed. Then, for each unique compound in NIST20 (identified by InChIKey), we chose its most frequent adduct form and performed subformula-based spectra merging for MS/MS collected from the same MS instrument but under different collision energies.

For NIST20 distribution analysis, we plotted the distributions of 15 MLR features relevant to DBE value, hydrogen/carbon ratio, and heteroatom/carbon ratio, removing extreme outliers in advance using Tukey’s fences^9^ (k = 3). The skew normal distribution was chosen to estimate the 15 distributions for its broad scalability (**Supplementary Table 15 & Supplementary Fig. 8**), where the parameter optimization was achieved via the Broyden–Fletcher–Goldfarb–Shanno (BFGS) algorithm^10^. Notably, we differentiated between (de)protonated fragment ions and radical fragment ions (also between non-radical neutral losses and radical neutral losses) during the distribution analysis; distribution results of positive and negative ion mode data were also separately generated. These skew normal distribution curves are then used to calculate MS/MS-related features in the MLR process, aiding the quantification of the performance of precursor formula candidates in explaining fragment ions and neutral losses in MS/MS spectra.

### Supplementary Note 9 | MS1 isotope simulation workflow.

To train the MLR model, which takes both MS1 isotope and MS/MS as input, training data containing experimental MS1 isotopic patterns and MS/MS spectra are needed. Due to the limited amount of publicly available MS data that meet the condition above, we performed the isotope simulation to generate experimental-like MS1 isotopic patterns for chemicals in NIST20. The simulation workflow was carefully designed based on the experimental MS1 isotope analysis of 314 chemical standards. Here, the isotope simulation was carried out considering the first four isotopic peaks (M + 0 to M + 3).

The isotope simulation proceeds as follows.

1) The theoretical isotopic pattern of the target chemical compound was computed via the open-source Chemistry Development Kit (CDK).
2) Given a summation window of width τ (mass resolution-dependent), we summed the isotope intensities for adjacent isotopologues within τ (e.g., ^13^CH_4_ & ^12^CH_3_D; ^13^CH_3_D & ^12^CH_2_D_2_. See **Supplementary Fig. 9**).
3) Isotope peak intensities were then normalized using the monoisotopic peak M + 0.
4) For isotopic peaks of M + 1 to M + 3, intensity shifting factors ($\alpha$) were sampled from a Gaussian distribution with a mean of 0. The standard deviation ($\sigma$) varied for different isotopic peaks.

$\alpha_{i}\sim N\left( 0,{\sigma_{i}}^{2} \right)$ where $i=1, 2, 3$

5) We then determined sign factors ($\beta$) for M + 1 to M + 3 isotopic peaks, where $\beta=\pm1$. For M + 1 isotopic peak, its sign factor $\beta_{1}$ has an equal chance to be 1 or −1. For M + 2 and M + 3 peaks, their sign factors $\beta_{i}$ ($i=2, 3$) rely highly on $\beta_{i-1}$ values.

$$P\left( \beta_{1}=1 \right)=P\left( \beta_{1}=-1 \right)=0.5$$

$P\left( \beta_{i}=\beta_{i-1} \right)=\rho$ where $i=2, 3$

6) The simulated intensities were calculated for the M + 1 to M + 3 isotopic peaks based on their theoretical intensities. If $\alpha_{i}\beta_{i}<-1$, repeat steps 4 to 6.

${Int}_{sim, i}=\left( 1+\alpha_{i}\beta_{i} \right)\times{Int}_{theo,i}$ where $i=1, 2, 3$

7) Isotope peak intensities were renormalized; ready for isotope similarity calculation.

In this study, we set the parameters as $\sigma_{1}$ = 0.11, $\sigma_{2}$ = 0.16, $\sigma_{3}$ = 0.19, and $\rho$ = 0.70. Of note, the simulation of mass deviations in isotopic peaks was not performed here. We argue that this simulation process is unnecessary due to the intrinsic advantage of the applied isotope similarity algorithm. The existence of the summation window enables the mass shifting of isotopic peaks within a reasonable range without affecting the final isotope similarity scores. However, the simulation of precursor mass deviations is essential in MLR training.

To validate the above isotope simulation workflow, we simulated the isotopic patterns of 314 chemical standards ten times in parallel. Isotope similarity scores were calculated between the simulated and theoretical isotopes, which we note here as “simulated pairs”. Compared to the isotope similarity scores of “correct pairs” (experimental & corresponding theoretical isotopes), their density plots were visually highly similar (**Supplementary Fig. 10**). Furthermore, statistical analysis was conducted between the isotope similarity score distributions of “simulated pairs” and “correct pairs”, and no statistical significance can be observed between their medians or empirical distribution functions (two-sided Mann-Whitney U test: *P* > 0.70; two-sided two-sample Kolmogorov–Smirnov test: *P* > 0.10). The above results indicate that our simulated isotopes are close to experimental conditions in terms of how close their isotope similarity scores are to theoretical. As such, we were able to train the MLR model using NIST20 spectra and their corresponding simulated MS1 isotopic patterns, which extensively expanded the scope of available training data.

### Supplementary Note 10 | Data augmentation in MLR training.

In the NIST20 database, there can be tens of available reference MS/MS spectra for a single chemical compound, collected in different ionization modes, adduct forms, collision energies, or even MS instruments. We merged MS/MS spectra for unique chemical compounds prior to MLR training for structure-disjoint evaluation. This led to higher fragment numbers in merged MS/MS than in experimental MS/MS. To this end, data augmentation is required for two purposes: 1) adjusting the fragment number of MS/MS spectra in training data closer to experimental cases and 2) increasing the generalization of trained models so that correct annotation results can be obtained even without many available fragments. The data augmentation workflow in the MLR training process was designed as follows.

For high-resolution MS/MS spectra, fragments in each merged MS/MS spectrum were first sorted by peak intensity. From a merged MS/MS with a fragment count of ***m***, we generated two augmented MS/MS spectra by reserving top $ceiling(\frac{\boldsymbol{m}}{2})$ and top $ceiling(\frac{\boldsymbol{m}}{5})$ fragments, where the function *ceiling* calculates the smallest integer no less than the numerical input. The amount of available training MS/MS was thus doubled, and their fragment counts were rectified closer to experimental cases.

For low-resolution MS/MS spectra, we modified the reserved fragment numbers as ***m*** and top $ceiling(\frac{4\boldsymbol{m}}{5})$, as low-resolution merged MS/MS have much fewer fragments than high-resolution merged MS/MS.

### Supplementary Note 11 | Evaluation metrics used in MLR training.

During the MLR training process, we used normalized discounted cumulative gain (NDCG) as the evaluation metric to optimize MLR models, which is calculated based on the value of discounted cumulative gain (DCG). The calculation formulae of DCG and NDCG are shown as below.

$$DCG@N=\sum_{i=1}^{N} \frac{g_{i}}{\ln\left( i+1 \right)}$$

$$NDCG@N=\frac{DCG@N}{MaxDCG@N}$$

where $g_{i}$ is the relevance gain at the *i*-th position in the ranking list (correct answers receive 1; incorrect answers receive 0); “$@N$” refers to the *N*-th element; MaxDCG refers to the maximum (ideal) DCG if all candidates are ranked in the ideal order.

Reference:

https://docs.microsoft.com/en-us/dotnet/api/microsoft.ml.data.rankingmetrics?view=ml-dotnet

### Supplementary Note 12 | Experiment-specific global peak annotation.

Experiment-specific global peak annotation is achieved by node scoring, edge scoring, and global optimization. We discuss these details below.

Node scoring reflects each formula candidate’s quality of explanation for the target peak. Here the node score is purely the calibrated posterior probability provided by Platt scaling. The posterior probability is an integrated numeric value derived from the MLR task that takes both MS1 and MS/MS information into account. Notably, we do not implement any bonus or penalty scores based on meta-scores, meaning that the formula occurrences in common chemical databases, such as HMDB or KEGG, are not included in the node scoring system at this stage. Although users can choose to incorporate meta-scores in bottom-up MS/MS interrogation, global peak annotation is entirely free of meta-scores, enabling the broad scalability of annotation in BUDDY for different sample types collected from various species.

Valid edges are constructed considering both biochemical and abiotic connections. For biochemical connections, the formula difference between two peaks should match a biotransformation reaction provided in the list (**Supplementary Table 12**). If MS/MS spectra are available for both peaks, a minimum MS/MS similarity score of 0.6 is required as the GNPS algorithm^11^ is adopted to encompass potential biochemical transformations between two metabolic features. For abiotic connections (adduct formation or in-source fragmentation), retention time match is a prerequisite. For the identification of in-source fragmentation, we ensure that the precursor mass of one feature appears as a fragment ion in the MS/MS spectrum of the other feature. Edge scores contain a basic score and MS/MS similarity score, meaning that feature connections with valid MS/MS matching are awarded.

We next set up an integer linear programming (ILP) problem. Linear constraints are made to ensure that: (1) there exists one and only one selected formula annotation for each peak (node), and (2) a peak connection (edge) only exists when corresponding formulae in both peaks are selected. The above constraints guarantee a self-consistent network. Finally, we define the goal function to be optimized as follows:

$$argmax(\sum_{i\in node} P_{i}+\sum_{k\in edge} (\alpha\times S_{k}+\beta))$$

where P_i_ is the Platt probability of the selected formula for each node, S_k_ is the MS/MS similarity score between two metabolic features. Notably, only valid edges representing biochemical or abiotic connections are included. Parameters α and β are included to adjust the scoring weight between node scores and edge scores. Here, we set α=0.10, β=0.05 for abiotic connections and β=0.002 for biochemical connections. Parameters are set up empirically without in-depth optimization. A slightly higher basic score (β) is given to abiotic connections owing to its high confidence and prevalence.

### Supplementary Note 13 | Parameter settings for evaluations on MS/MS libraries.

- Parameter settings in SIRIUS

For MS/MS reference spectra collected in QTOF instruments:

- Instrument: QTOF (MS1: 10 ppm)
- MS/MS MassDev: 20 ppm
- Possible ionization: [M + H]^+^ for positive ion mode, [M − H]^−^ for negative ion mode
- Tree timeout: 50
- Compound timeout: 200
- Chemical elements are set to CHNOPSFClBrI
- All other settings are default

For MS/MS reference spectra collected in Orbitrap instruments:

- Instrument: Orbitrap (MS1: 5 ppm)
- MS/MS MassDev: 10 ppm
- Possible ionization: [M + H]^+^ for positive ion mode, [M − H]^−^ for negative ion mode
- Tree timeout: 50
- Compound timeout: 200
- Chemical elements are set to CHNOPSFClBrI
- All other settings are default
- Parameter settings in BUDDY

For MS/MS reference spectra collected in QTOF instruments:

- Task: Bottom-up MS/MS interrogation
- MS1 tolerance: 10 ppm
- MS/MS tolerance: 20 ppm
- No formula database restriction implemented
- Chemical elements are set to CHNOPSFClBrI
- Timeout for a single query: 30 s
- All other settings are default

For MS/MS reference spectra collected in Orbitrap instruments:

- Task: Bottom-up MS/MS interrogation
- MS1 tolerance: 5 ppm
- MS/MS tolerance: 10 ppm
- No formula database restriction implemented
- Chemical elements are set to CHNOPSFClBrI
- Timeout for a single query: 30 s
- All other settings are default

### Supplementary Note 14 | Parameter settings in MS-DIAL for LC-MS/MS data preprocessing.

- Data collection
  - MS1 tolerance: 0.01 Da (Orbitrap); 0.02 Da (QTOF)
  - MS/MS tolerance: 0.025 Da (Orbitrap); 0.05 Da (QTOF)
- Peak detection
  - Minimum peak height: 1000 amplitude
  - Mass slice width: 0.025 Da (Orbitrap); 0.05 Da (QTOF)
- Adduct
  - Positive ion mode: [M + H]^+^, [M + NH_4_]^+^, [M + Na]^+^, [M + K]^+^, [M + H − H_2_O]^+^, [M + H − 2H_2_O]^+^, [2M + H]^+^, [2M + NH_4_]^+^
  - Negative ion mode: [M − H]^−^, [M − H_2_O − H]^−^, [M + Cl]^−^, [M + FA − H]^−^, [M + Hac − H]^−^, [2M − H]^−^
- Alignment
  - Retention time tolerance: 0.1 min
  - MS1 tolerance: 0.01 Da (Orbitrap); 0.02 Da (QTOF)

### Supplementary Note 15 | Parameter settings for evaluations on public LC-MS/MS datasets.

- Parameter settings in SIRIUS

For LC-MS/MS data collected in QTOF instruments:

- Instrument: QTOF (MS1: 10 ppm)
- MS/MS MassDev: 20 ppm
- Possible ionization: [M + H]^+^ for positive ion mode, [M − H]^−^ for negative ion mode
- Tree timeout: 50
- Compound timeout: 200
- Chemical elements are set to CHNOPSFClBrI
- All other settings are default

For LC-MS/MS data collected in Orbitrap instruments:

- Instrument: Orbitrap (MS1: 5 ppm)
- MS/MS MassDev: 10 ppm
- Possible ionization: [M + H]^+^ for positive ion mode, [M − H]^−^ for negative ion mode
- Tree timeout: 50
- Compound timeout: 200
- Chemical elements are set to CHNOPSFClBrI
- All other settings are default
- Parameter settings in BUDDY

For LC-MS/MS data collected in QTOF instruments:

- Task: Bottom-up MS/MS interrogation
- MS1 tolerance: 10 ppm
- MS/MS tolerance: 20 ppm
- No formula database restriction implemented
- Chemical elements are set to CHNOPSFClBrI
- Timeout for a single query: 30 s
- All other settings are default

For LC-MS/MS data collected in Orbitrap instruments:

- Task: Bottom-up MS/MS interrogation
- MS1 tolerance: 5 ppm
- MS/MS tolerance: 10 ppm
- No formula database restriction implemented
- Chemical elements are set to CHNOPSFClBrI
- Timeout for a single query: 30 s
- All other settings are default

### Supplementary Note 16 | Parameter settings for applying BUDDY on NIST human fecal material standards.

−	Task: MS/MS library search, Bottom-up MS/MS interrogation & Experiment-specific global peak annotation

−	MS1 tolerance: 5 ppm

−	MS/MS tolerance: 10 ppm

−	MS/MS library search

−	MS/MS reference database: NIST20

−	Searching algorithm: dot product

−	Identification threshold: Similarity score: 0.7; Matched fragment count: 6

−	No formula database restriction implemented

−	No elemental ratio restriction

−	All other settings are default

### Supplementary Note 17 | Parameter settings for applying BUDDY on ARUS MS/MS libraries.

−	Task: Bottom-up MS/MS interrogation

−	MS1 tolerance: 5 ppm

−	MS/MS tolerance: 10 ppm

−	No formula database restriction implemented

−	Chemical elements are set to CHNOPSFClBrI

−	Timeout for a single query: 30 s

−	All other settings are default

References

1.	Sud, M. et al. LMSD: LIPID MAPS structure database. *Nucleic Acids Research* **35**, D527-D532 (2007).

2.	Moorthy, A.S., Wallace, W.E., Kearsley, A.J., Tchekhovskoi, D.V. & Stein, S.E. Combining Fragment-Ion and Neutral-Loss Matching during Mass Spectral Library Searching: A New General Purpose Algorithm Applicable to Illicit Drug Identification. *Anal Chem* **89**, 13261-13268 (2017).

3.	Wang, M. et al. Mass spectrometry searches using MASST. *Nature Biotechnology* **38**, 23-26 (2020).

4.	Hofmann, A.F., Hagey, L.R. & Krasowski, M.D. Bile salts of vertebrates: structural variation and possible evolutionary significance[S]. *Journal of Lipid Research* **51**, 226-246 (2010).

5.	Yamaguchi, K., Miyaguchi, H., Ohno, Y. & Kanawaku, Y. Qualitative analysis of 7- and 8-hydroxyzolpidem and discovery of novel zolpidem metabolites in postmortem urine using liquid chromatography–tandem mass spectrometry. *Forensic Toxicology* (2022).

6.	Xing, S. & Huan, T. Radical fragment ions in collision-induced dissociation-based tandem mass spectrometry. *Analytica Chimica Acta* **1200**, 339613 (2022).

7.	Kind, T. & Fiehn, O. Seven Golden Rules for heuristic filtering of molecular formulas obtained by accurate mass spectrometry. *BMC Bioinformatics* **8**, 105 (2007).

8.	Xing, S. et al. Recognizing Contamination Fragment Ions in Liquid Chromatography–Tandem Mass Spectrometry Data. *Journal of the American Society for Mass Spectrometry* (2021).

9.	Tukey, J.W. Exploratory Data Analysis. (Addison-Wesley, Reading, Mass.; Menlo Park, Calif.; London; Amsterdam; 1977).

10.	Fletcher, R. Practical methods of optimization. (2013).

11.	Watrous, J. et al. Mass spectral molecular networking of living microbial colonies. *Proc Natl Acad Sci U S A* **109**, E1743-1752 (2012).
